# BUB1 regulates non-homologous end joining pathway to mediate radioresistance in triple-negative breast cancer

**DOI:** 10.1101/2024.05.07.592812

**Authors:** Sushmitha Sriramulu, Shivani Thoidingjam, Wei-Min Chen, Oudai Hassan, Farzan Siddiqui, Stephen L Brown, Benjamin Movsas, Michael D Green, Anthony J Davis, Corey Speers, Eleanor Walker, Shyam Nyati

## Abstract

**Background:** Triple-negative breast cancer (TNBC) is a highly aggressive form of breast cancer subtype often treated with radiotherapy (RT). Due to its intrinsic heterogeneity and lack of effective targets, it is crucial to identify novel molecular targets that would increase RT efficacy. Here we demonstrate the role of BUB1 (cell cycle Ser/Thr kinase) in TNBC radioresistance and offer a novel strategy to improve TNBC treatment.

**Methods:** Gene expression analysis was performed to look at genes upregulated in TNBC patient samples compared to other subtypes. Cell proliferation and clonogenic survivals assays determined the IC_50_ of BUB1 inhibitor (BAY1816032) and radiation enhancement ratio (rER) with pharmacologic and genomic BUB1 inhibition. Mammary fat pad xenografts experiments were performed in CB17/SCID. The mechanism through which BUB1 inhibitor sensitizes TNBC cells to radiotherapy was delineated by γ-H2AX foci assays, BLRR, Immunoblotting, qPCR, CHX chase, and cell fractionation assays.

**Results:** BUB1 is overexpressed in BC and its expression is considerably elevated in TNBC with poor survival outcomes. Pharmacological or genomic ablation of BUB1 sensitized multiple TNBC cell lines to cell killing by radiation, although breast epithelial cells showed no radiosensitization with BUB1 inhibition. Kinase function of BUB1 is mainly accountable for this radiosensitization phenotype. BUB1 ablation also led to radiosensitization in TNBC tumor xenografts with significantly increased tumor growth delay and overall survival. Mechanistically, BUB1 ablation inhibited the repair of radiation-induced DNA double strand breaks (DSBs). BUB1 ablation stabilized phospho-DNAPKcs (S2056) following RT such that half-lives could not be estimated. In contrast, RT alone caused BUB1 stabilization, but pre-treatment with BUB1 inhibitor prevented stabilization (t_1/2_, ∼8 h). Nuclear and chromatin-enriched fractionations illustrated an increase in recruitment of phospho– and total-DNAPK, and KAP1 to chromatin indicating that BUB1 is indispensable in the activation and recruitment of non-homologous end joining (NHEJ) proteins to DSBs. Additionally, BUB1 staining of TNBC tissue microarrays demonstrated significant correlation of BUB1 protein expression with tumor grade.

**Conclusions:** BUB1 ablation sensitizes TNBC cell lines and xenografts to RT and BUB1 mediated radiosensitization may occur through NHEJ. Together, these results highlight BUB1 as a novel molecular target for radiosensitization in women with TNBC.

## Background

Breast cancer (BC) affects more than 2 million women worldwide each year. Triple-negative breast cancer (TNBC) is the most lethal subtype of BC and while effective targeted therapies exist for the prevention and treatment of ER-positive breast cancer, no effective targeted therapy exists for TNBC. TNBC tend to be more aggressive, occur in younger women, and are less likely to be cured by adjuvant therapy (1). As radiotherapy is standard in the management of BC, there is a need to identify molecular targets with potential to increase the efficacy of radiation therapy (RT). To this end, DNA damage repair pathways are of interest.

DNA damage is a critical determinant of radiation-induced cell death (2). Radiation mediated base damages and single strand breaks (SSBs) are more efficiently repaired by cells, whereas double strand breaks (DSBs) are more difficult to repair and, if unrepaired, lead to lethality in cells. The ability of cells to recognize and respond to DSBs is fundamental in determining the sensitivity (or resistance) of cells to radiation (3). DSB repair is comprised of two major and mechanistically distinct processes: non-homologous end-joining (NHEJ) and homologous recombination (HR). NHEJ involves directly ligating two broken DNA ends and is initiated by binding of Ku70/Ku80 hetero dimers at DSB sites (4). Ku70/Ku80 localization recruits DNA-dependent protein kinase (DNAPKcs) to the DSB site, followed by Artemis-dependent end-processing, strand synthesis by DNA polymerase-beta (POLβ) and strand ligation by XRCC4, ligase IV, and XLF complex (5). HR on the other hand is initiated by lesion recognition by ATM and processing of DSB ends by MRN complex (Mre1—Rad50-Nbs1). 53BP1 protein may play a role in pathway choice between NHEJ and HR (6, 7). Target-based radiosensitization approaches increase radiotherapy efficiency by selectively sensitizing tumor tissue to ionizing radiation (8). Several new molecular targets are currently being evaluated in clinical trials to measure their radiation sensitization potential (9).

Following DNA damage, cell cycle checkpoints are activated to block cell cycle progression and prevent propagation of cells with damaged DNA. Both DNA damage repair and cell cycle checkpoints are positively regulated by several kinases, including BUB1 (Budding uninhibited by benzimidazoles-1). BUB1 is a serine/threonine kinase implicated in chromosomal segregation during mitosis. BUB1 regulates cell-cycle and is known to impact DNA damage signaling. However, it is still uncertain how BUB1 contributes to radioresistance in TNBC. BUB1 is known to localize near DSB sites where early DNA damage sensor proteins such as phosphorylated H2AX are also recruited (10). Moreover, BUB1 co-localizes with 53BP1 suggesting a role in NHEJ pathway (10). Knockdown of BUB1 results in prolonged γH2AX foci and comet tail formation as well as hypersensitivity in response to ionizing radiation (11). Increased expression of BUB1 is associated with resistance to DNA-damaging agents (i.e. radiotherapy and some chemotherapies) (12) and we have shown that BUB1 inhibition reduces invasion and migration in cancer cell lines(13) through direct interaction with TGFβ receptors (14, 15). Moreover, BUB1 regulates cell cycle through its roles in spindle assembly checkpoint and chromosome alignment (16–18).

Here, we demonstrate that BUB1 is overexpressed in TNBC, and that its overexpression correlates with poorer outcome and radiation resistance. Moreover, we confirm that pharmacological or genomic ablation of BUB1 is cytotoxic to TNBC cell lines and leads to radiation sensitization. BUB1 ablation delays DSB repair as evident by prolonged γH2AX foci and affects NHEJ as evaluated by bioluminescent DNA damage repair reporters (BLRR). BUB1 inhibition causes significant decrease in tumor volume when combined with radiation in SUM159 mammary fat pad tumor xenograft models and demonstrates significant reduction in tumor cell proliferation as evaluated by Ki67 immunostaining of tumor sections. Additionally, our mechanistic studies show that BUB1 mediates radioresistance through impacting chromatin localization of core NHEJ proteins and increasing radiation mediated DNAPKcs phosphorylation and stability. Overall, our results provide evidence that BUB1 mediated radiation resistance takes place through NHEJ, specifically by regulating chromatin binding of key proteins and that combining BUB1 ablation with radiation could be an effective approach for radiosensitization of TNBC.

## Methods

### Gene expression data

Normalized expression data for the cell lines were downloaded from the EMBL-EBI ArrayExpress website as described in the original publication (19). The Hatzis gene expression and survival data were downloaded from the Gene Expression Omnibus (GEO) database with series number GSE25066 (20). A log-rank (Mantel-Cox) test was used for survival curve analyses. Data for the TCGA cohort was downloaded from http://tcga-data.nci.nih.gov. Expression levels were log transformed, median centered and scaled, subtype calls were based on previous description (21).

A receiver operating characteristic curve (ROC) was generated as an alternate way to measure the performance of BUB1 as a biomarker using area under the curve (AUC) as a metric, with an AUC >0.65 being considered of significant clinical value. BUB1 expression was evaluated as a continuous variable. BUB1 expression was measured by using RNA isolated from patients tumors at time of surgical expression, then log transformed values from the Affymetrix Human Genome U133A Array were assessed. Other clinical covariates included ER, PR, overall stage, size, nodal status, and PAM50 classification (p =0.0003).

### Gene expression and metastasis correlation

In vivo screening for metastases was performed using Chick Chorioallantoic Membrane assays in 21 preclinical breast cancer models with data published previously (22). Correlation coefficients were calculated using Pearson’s correlation methods.

### Cell culture

Triple-negative breast cancer cell lines (MDA-MB-231, MDA-MB-468, BT-549), normal breast epithelial cell line (MCF10A) and Estrogen Receptor (+), Progesterone Receptor (+) breast cancer cell line T47D were obtained from the American Type Culture Collection (ATCC). SUM159 cells were originally sourced from Steve P. Either (University of Michigan) and were acquired from Sofia Merajver (University of Michigan). SUM159 cells were grown in HAM’S F-12 media (Catalog No. 31765035, Thermo Fisher Scientific) supplemented with 5% FBS, 10 mM HEPES, 1 μg/ml Hydrocortisone, 6 μg/ml Insulin, and 1% Penicillin-Streptomycin. MDA-MB-231 and MDA-MB-468 cells were grown in DMEM media (Catalog No. 30-2002, ATCC) supplemented with 10% FBS and 1% Penicillin-Streptomycin. BT-549 and T-47D cells were grown in RPMI-1640 media (Catalog No. 30-2001, ATCC) supplemented with 10% FBS, 0.023 U/ml insulin, and 1% Penicillin-Streptomycin. MCF10A cells were also grown in RPMI-1640 media supplemented with 10% FBS, and 1% Penicillin-Streptomycin. All cell lines were maintained at 37LC in a 5% CO_2_ incubator and passaged at 70% confluence. Cell lines were routinely tested for mycoplasma contamination. Mutations in key genes is listed in **Supplementary Table S1** (See Additional file 1 for Supplementary figures and tables).

### Drug treatment and irradiation

A BUB1 inhibitor (BUB1i) BAY1816032 (Catalog No. HY-103020, MedChemExpress) and DNAPK inhibitor (DNAPKi) NU7441 (Catalog No. S2638, Selleckchem) were dissolved in DMSO (20 mM BUB1i and 15 mM DNAPKi) and stored at –80LC. For each experiment, a fresh vial was thawed, and any remaining stock solution was discarded. Working concentrations were made in media with serum and supplements and cells were exposed to a range of concentrations, from 125 nM to 1000 nM. Irradiation was performed 1 h after the drug treatment using a CIX-3 orthovoltage unit (Xstrahl Life Sciences) operating at 320 kV and 10 mA with 1 mm Cu filter.

### Proliferation assay

To investigate the effect of BAY1816032 on cell proliferation in TNBC cell lines, 2 x 10^3^ cells were plated into a 96-well plate 24 h prior to treatment. Cells were exposed to different concentrations of BUB1 inhibitor (BUB1i) ranging from 1 nM to 10 μM and cultured for 72 h. Cell proliferation was measured using alamarBlue (Catalog No. DAL1025, Thermo Fisher Scientific) following the manufacturer’s protocols. Absorbance was read at 570 nM on Synergy H1 Hybrid Reader (BioTek Instruments). Values were normalized to mock (DMSO/vehicle) treated cells. The IC_50_ values were estimated on GraphPad Prism (V9) using a non-linear regression best-fit equation.

### Clonogenic survival assay

Cells were plated in 6-well plates at different cell densities overnight. The next morning, cells were treated with BAY1816032 (125 nM to 1000 nM) for 1 hr and irradiated (2 to 6 Gy). Cells were allowed to grow for 7-23 days until visible colonies formed before being fixed and stained with methanol and crystal violet. All the colonies with >50 cells were manually counted, and the cell survival was plotted using GraphPad (V9). Plating efficiency (PE %) was estimated as: (100 x Number of colonies formed / Number of cells plated x 100). Radiation enhancement ratios (rER) were determined from the survival curve using the formula: D bar of varying inhibitor concentrations / D bar of vehicle (DMSO) (Microsoft Excel) which indicates radiation dose to produce some level of cell killing in the absence of inhibitor (i.e., vehicle) divided by the radiation dose in the presence of the inhibitor to produce the same level of cell kill. rER >1 was considered to be radiation sensitization while rER <1 was radiation resistance/protection.

### Transfections

Cells were seeded in 6-well plates overnight and the transfection was performed with 60% confluent cells. The siGENOME SMARTPool siRNA for human BUB1 and DNAPK (gene ID: PRKDC) were purchased from Dharmacon. Next morning, 100 nM siRNAs were diluted in Opti-MEM reduced serum media (Catalog No. 31985062, Thermo Fisher Scientific) and transfected using Lipofectamine RNAiMAX (Catalog No. 13778075, Thermo Fisher Scientific). Diluted siRNAs and lipofectamine reagent were separately incubated for 5 mins at RT before being combined and incubated further for 20-30 mins after combining them. The siRNA-lipid complex was added to the cells in plain media without serum and antibiotics. After 48 h of transfection, the transfected cells were used for further experiments. BUB1 Wild-type (WT) and Kinase-dead (KD) plasmids were a kind gift from Dr. Hongtao Yu (UT Southwestern). The BUB1 plasmids or siRNA with plasmids were transfected with Lipofectamine 2000 (Catalog No. 11668500, Thermo Fisher Scientific) per manufacture’s protocol.

### Generation of BUB1 CRISPR knockout cell lines

Cells were transfected with CRISPR/CAS9 ribonucleoprotein (RNP) for generating BUB1 knockout cell lines. BUB1 sgRNAs were designed with the CRISPR tool (www.benchling.com) and synthesized by Integrated DNA Technologies (IDT). We combined two sgRNAs (sgRNA1 and sgrna2) in this experiment each targeted different exons (exon 2 and 3) for better knockout efficiency. Purified CAS9 protein was purchased from IDT (Catalog No. 1081058) while Lipofectamine RNAiMAX (Catalog No. 11668027) was from Thermo Fisher Scientific. TNBC cell lines were plated in 2 wells of a 24-well plate overnight. Next morning, the cells in one well were transfected by combining BUB1 sgRNA1 and sgRNA2 (each 300 ng/well), Cas9 protein (1 µg/well) using Lipofectamine RNAiMAX (3 µl/well) and the other well was used as a negative control without sgRNAs. 24-hours after transfection, cells were trypsinized and plated in 96-well plate at 1 cell/well. Cells were allowed to grow until colonies formed (2-4 weeks) and expanded into 24 well plates. Genomic DNA was isolated from these clones using the QuickExtract DNA extraction solution (Catalog No. QE09050, Lucigen) following manufacturer’s protocol. The extracted DNA was PCR-amplified with following conditions: 98LC for 30 s, 98LC for 10 s, 61.5LC for 30 s, 72LC for 23 s, and 72LC for 10 mins for 34 cycles. The putative BUB1 null clones were sequence verified (Sanger sequencing, Azenta Life Sciences, NJ, USA) and absence of BUB1 protein was confirmed by western immunoblotting. The efficiency of BUB1 CRISPR knockout was estimated by Synthego ICE software. gRNA sequences for BUB1 knockout, primer sequences for PCR amplification and Sanger sequencing are listed in **Supplementary Table S2**.

### Immunoblotting

Total protein was extracted using IP-lysis buffer (50mM Tris PH 7.4, 1% NP40, 0.25% Deoxycholate sodium salt, 150mM NaCl, 10% Glycerol, and 1mM EDTA) supplemented with PhosStop (Roche), Protease inhibitor (Roche), Sodium Ortho Vanadate, Sodium fluoride, PMSF, and β-Glycerol phosphate (2 µM each). Protein concentrations were determined using Pierce BCA protein assay kit (Catalog No. 23225, Thermo Fisher Scientific) and equal amounts of samples were loaded on NuPAGE 4-12%, Bis-Tris Midi protein gels (Catalog No. WG1402BOX, Thermo Fisher Scientific) along with SeeBlue Plus2 Pre-stained Protein Standard (Catalog No. LC5925, Thermo Fisher Scientific). Samples were transferred to Immobilon-P PVDF membranes (Catalog No. IPVH00010, Millipore). The blots were blocked using 5% non-fat dry milk (Catalog No. 1706404, BioRad) and/or 5% BSA and incubated with primary antibodies at 4□C overnight. Membranes were incubated with HRP-tagged secondary antibodies and protein bands were detected using ECL Prime western blotting system (Catalog No. GERPN2232, Millipore Sigma). Protein band density was measured using ImageJ 1.52a. Specific antibody information and dilutions are listed in **Supplementary Table S3**.

### Animal studies

Fox Chase SCID female mice (CB17/lcr Prkdcscid/lcrlcoCrl; 8 weeks old) (N = 52) were procured from Charles River Laboratories through the Department of Bioresources, Henry Ford Health. Mice were acclimatized for a week and housed at the Animal Facility, E&R building, Henry Ford Hospital. Experimental animals were housed and handled in accordance with protocols approved by IACUC of Henry Ford Health (protocol # 00001298). We used >9-10 mice per treatment group (2 tumors/mouse). After injecting SUM159 cells (1 x 10^6^ bilaterally) into the 4^th^ mammary fat pads, animals were randomly assigned to receive treatment once the tumors reached a size of about 80 mm^3^. BAY1816032 (25 mg/kg, *in vivo* grade, Catalog No. CT-BAY181, Chemietek) dissolved in 50% PEG 400, 10% DMSO, and 40% saline was given orally twice daily (5 days) for four weeks. RT was administered in three 5 Gy fractions over 5 days (total 15 Gy) using the small animal radiation research platform (SARRP, Xstrahl Life Sciences). Animals wherein tumors were generated with SUM159 BUB1 CRISPR KO cells were treated only with radiation or sham irradiated. Tumor volume and animal body weights were measured twice a week using a digital vernier caliper and tumor volume was calculated using the formula: (Length x Width^2^) x 3.14/6. When the tumor volume reached >1000 mm^3^, mice were euthanized according to IACUC guidelines. Linear mixed model (LMM) of log2 (tumor volume) was built on time and time* arm interaction. LMM was clustered by each tumor and nested within each mouse. 95% CI was 0.1 while p-value <0.001 was considered significant. Animal survival was estimated and depicted in a Kaplan-Meier survival plot. Logrank test were performed to estimate if the arms (treatment groups) were different (p<0.0001). Cox proportional hazards model with Firth’s penalized maximum likelihood bias reduction method was used to compare if experimental conditions resulted in significant differences.

### Immunohistochemical staining

Five random tumors from each treatment groups were harvested, fixed in buffered formalin and paraffin embedded. Histological sections from individual paraffin-embedded xenograft tumor tissues were initially deparaffinized and rehydrated. These tumor sections were stained at the Histology core (Henry Ford Health) according to the manufacturer’s protocols (Ki-67 IHC MIB-1, Dako Omnis). Proliferating cells were immunostained with FLEX monoclonal mouse anti-human Ki-67 (Catalog No. GA626, Ready-to-use (Dako Omnis), Clone MIB-1, Agilent). Images of the microscopic slides were taken under the light microscope at 20x magnification in two to three random fields for each tumor (N = 5, each arm). The % of Ki-67 positive cells was calculated using the formula: 100 x number of Ki-67 positive cells in treated / sham. H&E staining was also performed to assess the structural changes in the tumor sections.

### **γ**H2AX foci formation assay

Cells (1 x 10^5^) were plated into a 6-well plate containing glass coverslips (12 mm Catalog No. 633029, Carolina). After treatment with BUB1i or DNAPKi (1 μM) for an hour, cells were irradiated (4 Gy) and coverslips were collected at different time points. Coverslips were washed with ice-cold PBS, fixed with 2% w/v Sucrose, 0.2% Triton X-100 in formaldehyde, and permeabilized 0.5% triton in PBS for 10 min. Coverslips were blocked for 30 min at 4□C (2.5% Horse serum, 2.5% FBS, 0.5% w/v BSA, 0.05% Triton X-100 in PBS) and were incubated with Anti-phospho-Histone H2A.X (Ser139); Millipore) for 1 h at room temperature. After washing, coverslips were incubated with Goat anti-Mouse-Alexa Fluor 488 (Thermo Fisher Scientific) for 30 min at 4□C. DAPI (1 µg/ml) was used as a nuclear counter stain and the coverslips were mounted using ProLong Gold Antifade Mountant (Catalog No. P10144, Thermo Fisher Scientific) onto glass slides and observed under a microscope (Zeiss Axio Imager 2). At least 3 random fields were imaged for each condition. Cells with more than 10 γH2AX foci were scored as positive. The percentage of γH2AX foci positive cells was calculated as: (100 x number of γH2AX foci positive cells / Total number of cells counted).

### Bioluminescent NHEJ and HR reporter assays

The rate of DNA repair by NHEJ and HR was measured by the bioluminescent repair reporter (BLRR) kindly gifted by Dr. Christian E Badr, Harvard Medical School (23). Cells (2.5 x 10^5^) were plated in a 6-well plate and transfected with pLenti-BLRR (Addgene # 158958), pLenti-trGluc (Addgene # 158959), and pX330-gRNA (Addgene # 158973) plasmids with Lipofectamine 2000. After 48 h of transfection, the cells were reseeded into a 96-well plate, treated with BUB1i or DNAPKi (1 μM) for 1 h followed by RT (4 Gy), and replaced with fresh media. After 48 h of treatment, the cell supernatant was collected, centrifuged and 20 μl was transferred in a white opaque 96-well plate (Catalog No. IP-DP35F-96-W, Stellar Scientific). 1 mM Coelenterazine (Catalog No. 16123, Cayman Chemical) diluted to 80 μl was added to the supernatant and *Gaussia* luciferase activity (GLuc; HR efficiency) was measured for 0.8 s on Synergy H1 Hybrid (Biotek Instruments). 6.16 mM Vargulin (Catalog No. 305, NanoLight Technology) diluted in 50 μl was added to measure *Cypridina* luciferase activity (VLuc; NHEJ efficiency) with integration time of 1 s.

### Quantitative PCR

Cells (1.5 x 10^5^) were seeded in 6-well plates 24 h prior to treatment with BUB1i or DNAPKi (1 μM) and irradiation (4 Gy). Cells were harvested after 72 h and stored at –80□C. Total RNA was isolated with TRIzol (Catalog No. 15596026, Thermo Fisher Scientific) and concentration was measured on Nanodrop (Nanodrop 2000c, Thermo Fisher Scientific). RNA was reverse transcribed into cDNA using Super Script III Reverse Transcriptase kit (Catalog No. 18080044, Thermo Fisher Scientific), dNTPs (Catalog No. R0191, Thermo Fisher Scientific), and Random Primers (Catalog No. 48190011, Thermo Fisher Scientific). The qPCR was performed using Takyon Low ROX SYBR 2X MasterMix (Catalog No. UF-LSMT-B0701, Eurogentec) and *KiCqStart* pre-designed SYBR green gene-specific primers (**Supplementary Table S4**) in QuantStudio 6 Flex Real-Time PCR System (Applied Biosystems). Expression level for each gene was normalized to GAPDH for each experiment. All QRTPCR reactions were performed in triplicates and all experiments were repeated at least three times.

### Cycloheximide-chase assay

1.5 x 10^5^ cells were seeded in a 6-well plate over-night. Next morning cells were treated with 50 μM Cycloheximide (CHX; Catalog No. 14126, Cayman Chemical) to block nascent protein synthesis followed by BAY1816032 (1 μM) and 4 Gy irradiation. BUB1 CRISPR KO cells were treated with CHX followed by 4 Gy irradiation. Proteins were eluted at different time points (0 – 24h) by direct lysis (IP lysis buffer with 1.25X SDS protein loading buffer), sonicated, boiled for 7-8 mins before loading on the SDS-PAGE gels. Protein band density was quantified using ImageJ 1.52a software and calculated fold change using Microsoft Excel. The graphs were plotted in GraphPad Prism 9 software and the average half-life of BUB1 protein (t_1/2_) was determined using Microsoft Excel.

### Subcellular fractionation

To investigate the effect of BUB1 ablation on localization and movement of key DNA repair proteins on break sites and chromatin was investigated by subcellular fractionation assays. Different protein fractions were collected using Subcellular Protein Fractionation Kit (Catalog No. 78840, Thermo Fisher Scientific) according to the manufacturer’s protocols. Briefly, 2.5 x 10^6^ cells were plated in 100 mm petri dishes 48 h prior to treatment. BUB1i was added 1 h prior to irradiation (8 Gy) and allowed to recover for 10 mins. Cells were harvested, protein fractions were eluted as recommended and 40 μg protein was loaded onto NuPAGE 4-12%, Bis-Tris gels for western blot analysis.

### Laser micro-irradiation

U2-OS cells expressing YFP-tagged Ku80 and YFP-tagged DNA-PKcs were generated in the earlier studies (PMID: 22179609, PMID: 35580045). YFP-Ku80 and YFP-DNA-PKcs were transfected into U2-OS DNA-PKcs +/+ and −/− cells with JetPrime® (Polyplus transfection reagent, Catalog No. 101000027) following the manufacturer’s instructions. To observe the role of BUB1 in the accumulation of DNA-PKcs and KU80 at DNA DSBs, BUB1 was inhibited, and the cells were subjected to laser micro-irradiation. Twenty-four hours after the transfection, laser micro-irradiation and real-time recruitment were carried out using a Carl Zeiss Axiovert 200M microscope with a Plan-Apochromat 63X/NA 1.40 oil immersion objective (Carl Zeiss) as described in previous studies (24). A 365-nm pulsed nitrogen laser (Spectra-Physics) connected directly to the microscope’s epifluorescence path was used to create DSBs (24). During micro-irradiation, the cells were kept in an Invitrogen CO_2_-independent medium at 37°C. The fluorescence intensities of the micro-irradiated and control areas were measured using the Carl Zeiss Axiovision software, version 4.5. The irradiated area’s intensity was then normalized to the non-irradiated control area in accordance with earlier descriptions (25, 26).

### Tissue Microarray

Tissue microarray (TMA) panels of human breast carcinoma with adjacent normal breast tissues (BC081120f – 110 cores/110 cases and BR1191 – 119 cores/119 cases) were purchased from Tissue Array (formerly US Biomax, Derwood, MD). Breast TMAs were stained by the Histology Core-HFH with an anti-BUB1 antibody, (Catalog # ab195268, Clone EPR18947, Abcam, 1:50 dilution) following standard protocols. The slides were scanned/imaged using Aperio digital pathology slide scanner (Leica Biosystems). The TMAs were reviewed (manual scoring) by a blinded pathologist who provided the score of 0, 1+, 2+, 3+ that measures the staining intensity of BUB1, and the percentage of cells stained positive for BUB1. Graphs were plotted based on the staining intensity and % of cells positive for BUB1 to compare between normal and breast cancer tissues, molecular subtypes, tumor grades and stages.

### Statistical analysis

For the analyses of *in vitro* data, the Student’s t-test method was used in GraphPad Prism 9 software. Results are presented as mean ± standard error of the Mean (SEMs). All experiments were performed in triplicates and were repeated at least three times. Correlation coefficients were calculated using Pearson’s correlation methods. P < 0.05 was considered statistically significant. The statistical analysis of *in vivo* tumor growth data is presented under that section.

## Results

### BUB1 is overexpressed in TNBC and correlates with poorer survival and metastatic potential

In an effort to identify novel therapeutic targets for radiosensitization, we performed a screen focused on the human kinome to identify kinases upregulated in across 21 breast cancer cell lines that also impacted radiation sensitivity in human breast tumors (27). We identified a list of 52 kinases whose expression was significantly elevated in triple-negative breast cancer. We hypothesized that many of these kinases would govern mitogenic, metastatic, survival, or growth regulatory pathways critical to the development and dissemination of triple-negative breast cancer that could be readily targeted for the treatment of patients with triple-negative and basal-like breast cancer. To further characterize which of these 52 kinases played an important role in the aggressive features of triple-negative breast cancer, we combined expression, phenotypic, and clinical outcomes data to prioritize kinases that warranted further interrogation.

We prioritized those kinases that had the highest level of differential expression in triple-negative breast cancer, showed limited to no expression in normal tissues (including the mammary gland, thus were specific for breast cancer), were associated with clinically relevant outcomes, and for which we would be able to obtain or generate a specific inhibitor that was of clinical-grade quality to aid in translational efforts. To that end, BUB1 was one of the top nominated of the 52 kinases as it showed significantly elevated expression in triple-negative and basal-like breast cancers and limited expression in normal tissues (**Fig. 1A-B**, data based on expression in over 1000 patient tumors from TCGA). Additionally, BUB1 expression is significantly associated with basal-like and luminal B tumors and in triple-negative breast cancers (**Fig. 1C-D**). BUB1 expression is also much higher in breast cancer cell lines with basal-like characteristics and in cell lines with increased metastatic potential (**Fig. 1E**) (19). To further investigate the association of BUB1 expression with the metastatic potential of various breast cancer cell lines, we performed chick chorioallantoic membrane (CAM) assays on 21 breast cancer cell lines and quantitated the number of metastatic cells in the lungs and liver of chick embryos after injection of each of these 21 cell lines. This data was then correlated with BUB1 expression and there was a significant association between BUB1 expression and metastatic potential in this in vivo system (**Fig. 1F**, R^2^=0.64, p-value 0.004).

**Fig. 1.**
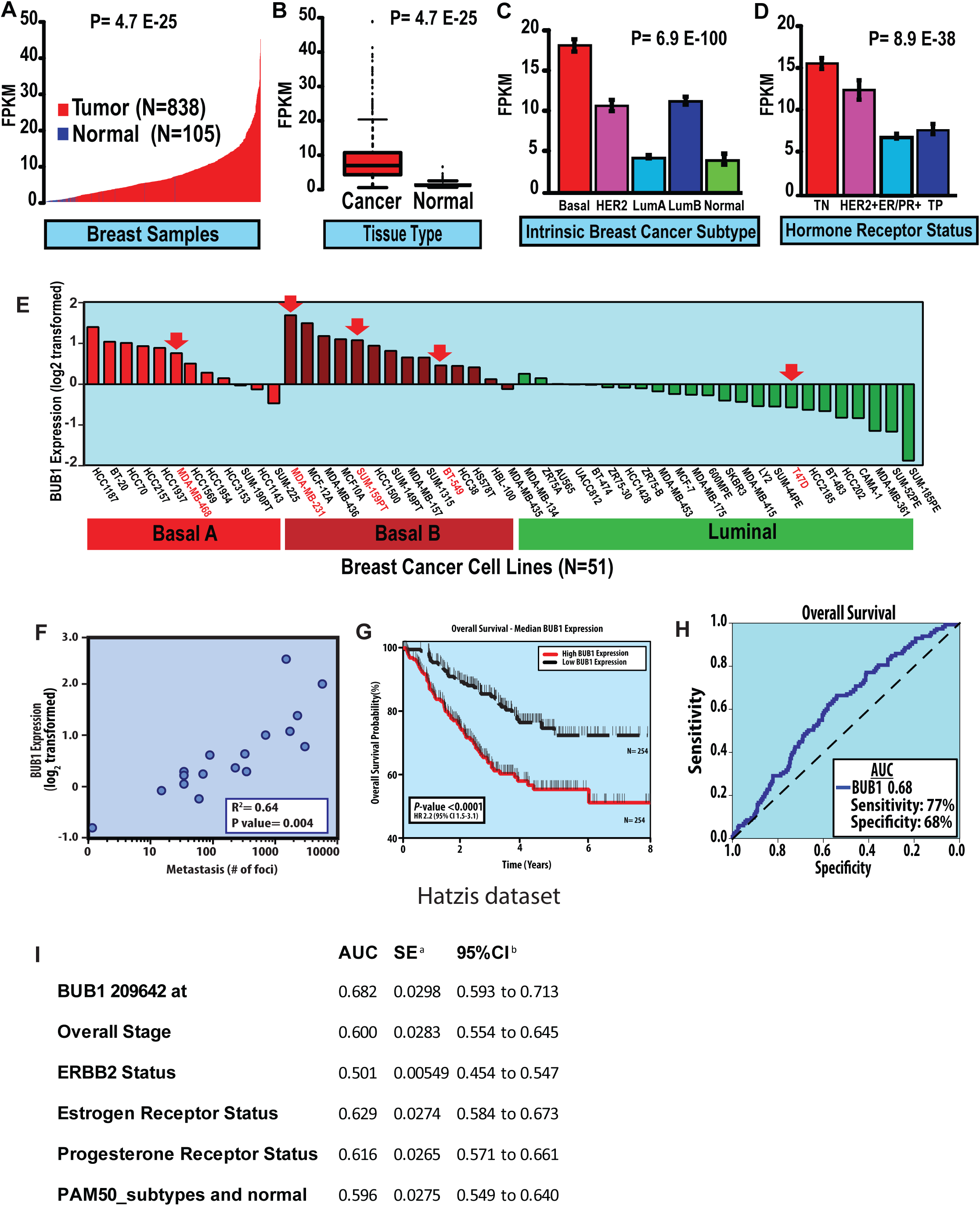
BUB1 is highly expressed in breast cancer compared to normal, non-malignant breast tissue and is associated with triple-negative and basal-like breast cancers. (A-B), BUB1 expression is significantly increased in breast tumors compared to normal breast tissue. (C), BUB1 expression is strongly associated with the PAM50-defined basal-like subtype of breast cancer and (D), is also significantly elevated in TNBC. (E), BUB1 expression is significantly increased in basal-like breast cancer cell lines. (F), BUB1 expression strongly correlates with metastatic potential to the lungs and liver as measured by CAM assay *in vivo*. All CAM assays performed at least in triplicate. (G), Kaplan-Meier survival plot demonstrate that high BUB1 levels are associated with worse overall survival in breast cancer patients (data from Hatzis et al, JAMA 2011). (H), On multivariable analysis, BUB1 expression discriminates overall survival with high sensitivity and specificity (AUC: 0.68, <0.01). (I), Raw data that was used for the analysis of the receiver operating characteristic curve (ROC). z statistic 3.631, ***P= 0.0003, ^a^DeLong et al., 1988 (70), ^b^Binomial exact

To investigate the clinical relevance of our findings, we assessed the impact of BUB1 expression on clinical outcomes. We found that BUB1 expression was significantly associated with poor outcomes (including higher mortality and increased rates of recurrence) in both women treated with chemotherapy and radiation therapy, the two most common adjuvant treatment modalities for women with breast cancer, with high BUB1 expression being strongly associated with worse overall survival in women with breast cancer (**Fig. 1G**) (20, 28). Furthermore, we demonstrate that BUB1 outperforms every other clinical or pathologic parameter (i.e., T-stage, grade, age, nodal status, ER, PR, Her2, margin, etc.) as a predictive biomarker of response (as measured by metastasis-free survival) to chemotherapy in a dataset of patients treated with paclitaxel and anthracycline-based chemotherapy with an AUC of 0.68 (**Fig. 1H**). The control group, tissue sample type, and detection method for Fig. 1H are described in the **Fig. 1I**.

### Pharmacological inhibition of BUB1 reduces viability of breast cancer cells

To study the effect of BUB1 inhibition in TNBC, we used the selective inhibitor of BUB1 kinase, BAY1816032. We assessed the effects of BAY1816032 on proliferation of TNBC (SUM159, MDA-MB-231, MDA-MB-468, BT-549) cells (**Fig. 2A-D**), luminal A subtype (T-47D) (**Fig. 2E**) and the non-tumorigenic human breast epithelial cell line, MCF10A (**Fig. 2F**). BAY1816032 is cytotoxic in all breast cancer cell lines tested with IC_50_ values ranging from 1.6 μM to 3.9 μM. However, BAY1816032 had less cell killing and/or growth inhibitory effects in MCF10A with IC_50_ around 18 μM. This response correlated with differential BUB1 mRNA expression (**Fig. 1E**) and BUB1 protein expression (**Fig 2G**). Based on these observations, we hypothesize that breast cancer cell lines that express high BUB1 would be radiosensitized by BAY1816032 while the cell lines that express low to moderate BUB1 would not.

**Fig. 2.**
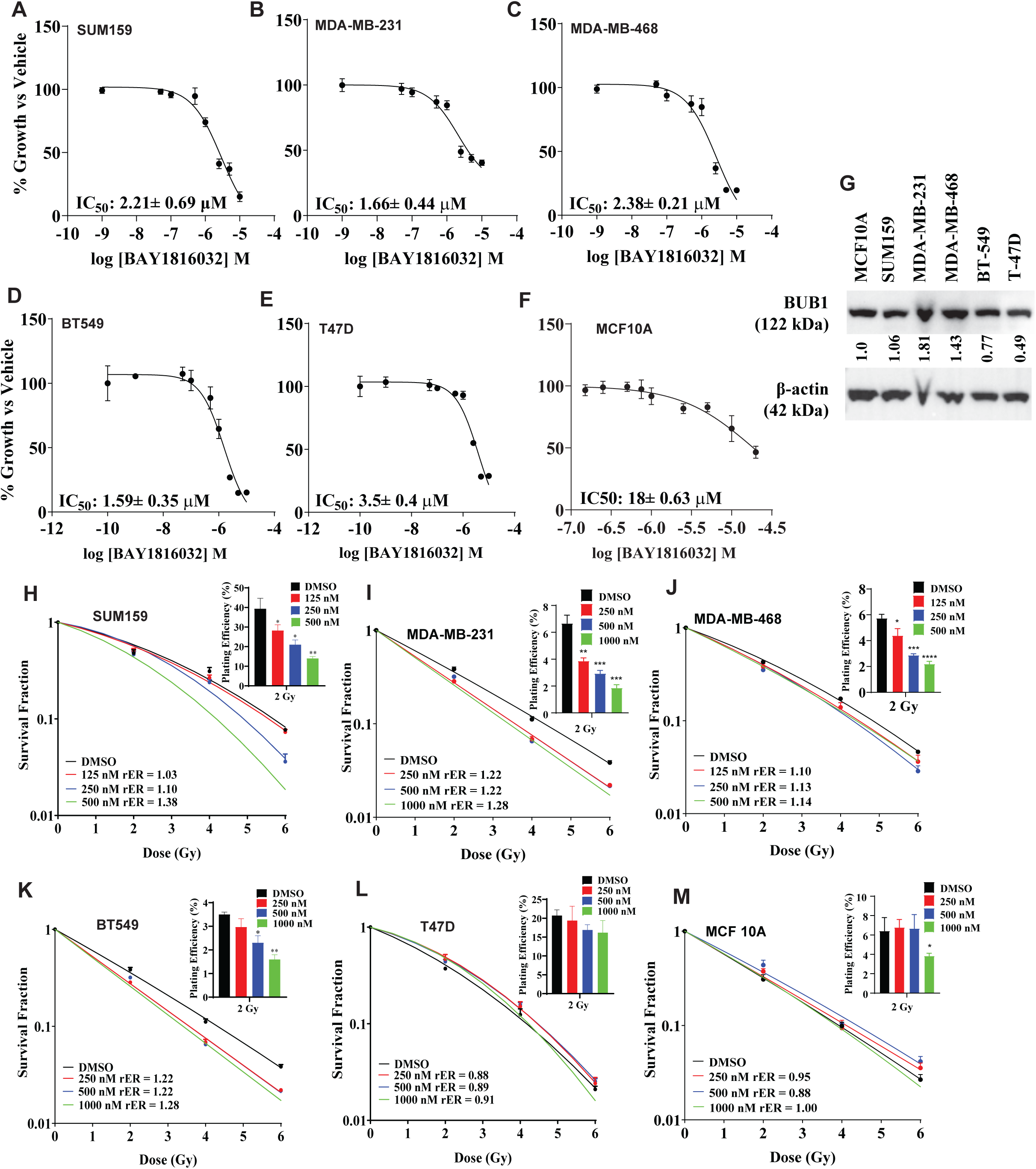
Effect of BUB1 inhibitor on cell proliferation in TNBC cell lines. BAY1816032 is cytotoxic to cells at low micromolar range (A) SUM159, IC_50_: 2.90 μM; (B) MDA-MB-231, IC_50_: 2.10 μM; (C) MDA-MB-468, IC_50_: 2.59 μM; (D) BT-549 IC_50_: 1.59 μM; (E) T-47D, IC_50_: 3.9μM; (F) MCF10A, IC_50_: 18 μM. (G) BUB1 protein expression in cell lines by immunoblotting; Pharmacological inhibition of BUB1 induces radiosensitivity in TNBC cell lines: (H) SUM159, (I) MDA-MB-231, (J) MDA-MB-468, (K) BT-549, (L) T-47D, and (M) MCF10A. P-values were defined as * P≤0.05, ** P≤0.01, *** P≤0.001, **** P≤0.0001.

### BUB1 inhibition causes durable radiosensitization in TNBC cell lines

We evaluated the effect of BAY1816032 on radiation sensitivity in Basal A (MDA-MB-468), Basal B (MDA-MB-231, SUM159, BT-549) (**Fig. 2H-K**) and Luminal A (T-47D) (**Fig. 2L**) cell lines by clonogenic survival assays. High levels of BUB1 expressed in selected Basal A and B cell lines (*BUB1-high*) while expressed at low level in Luminal cells (*BUB1-low*). *BUB1-high* cells were radiosensitized by BAY1816032 (rER from 1.1 to 1.38) while the radiation sensitivity of *BUB1-low* cells did not increase with BAY1816032 (rER 0.91). As expected, BAY1816032 had no effect on radiosensitivity in MCF10A cells (**Fig. 2M**). BAY1816032 led to a significant dose-dependent reduction in the surviving fraction at 2 Gy (SF-2 Gy) in *BUB1-high* cells indicating that BUB1 kinase function is important for radioresistance. Moreover, BAY1816032 did not significantly impacted SF-2Gy in *BUB1-low* cells.

### Genomic depletion of BUB1 is cytotoxic and makes TNBC cells radiosensitive

We evaluated the effect of BUB1 genomic depletion on cell survival and radiation sensitivity. SUM159 and MDA-MB-231 cells were transiently transfected with an increasing concentration of BUB1 siRNA (20, 60 and 100 nM) or control siRNA (100 nM) and cell viability was measured by alamarBlue assay (**Fig. 3A-B**). The siRNA-mediated BUB1 depletion demonstrated a dose-dependency on cell survival. Additionally, BUB1 was depleted in MDA-MB-468, BT-549 and T-47D cells which also exhibited significant reduction in cell viability as compared to control siRNA (**Fig. 3C-E**). DNAPKcs (gene ID: 5591, *PRKDC*) siRNA was used as a positive control since its inhibition or knockdown is known to reduce cell survival(29) because of the role it plays in DNA DSB repair process (30).

**Fig. 3.**
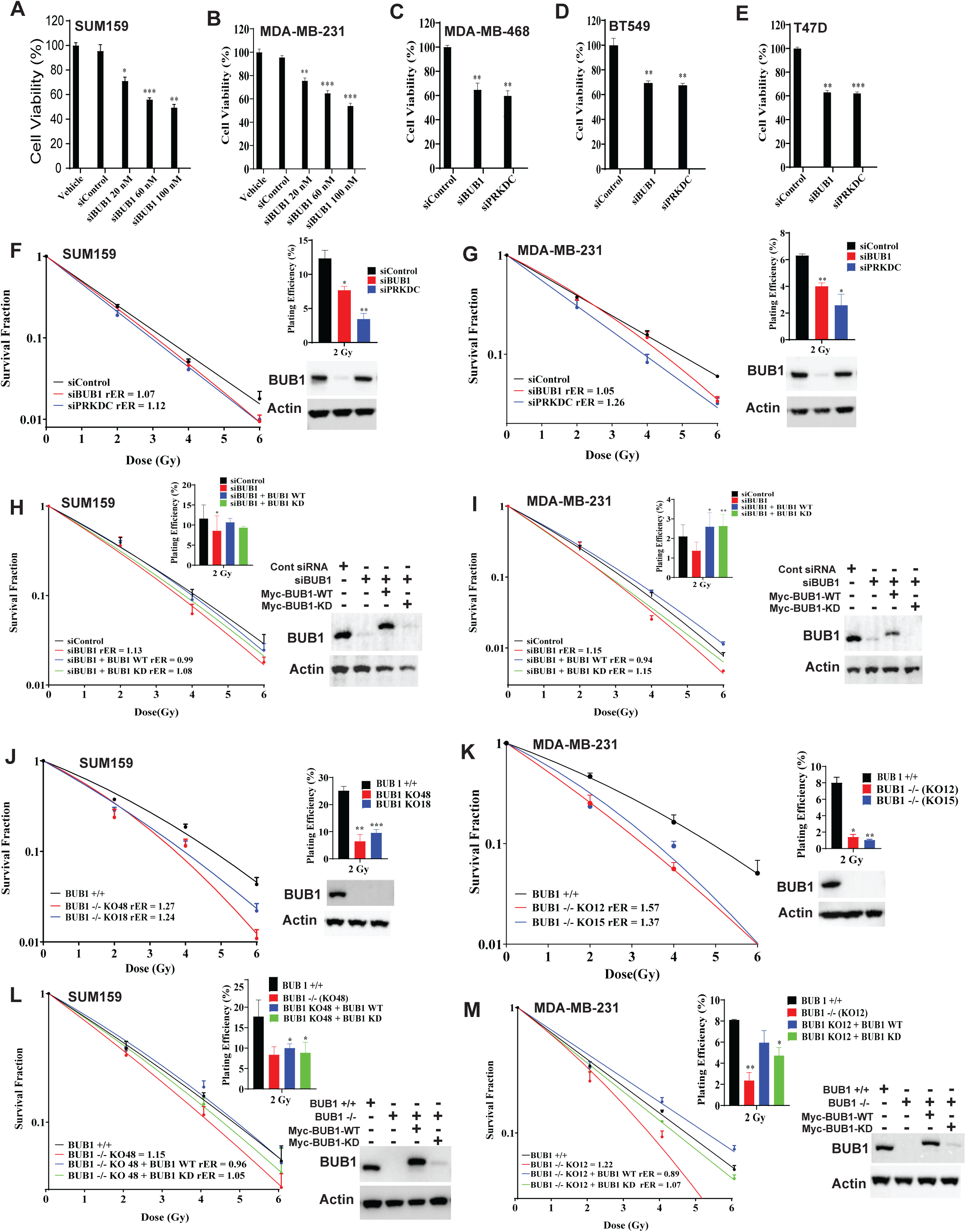
Effect of BUB1 genomic depletion on cell survival and radiation sensitivity. Transient transfection of BUB1 siRNA (20, 60 and 100 nM) or control siRNA (100 nM) measured cell viability using alamarBlue assay in (A) SUM159, (B) MDA-MB-231, (C) MDA-MB-468, (D) BT-549, and (E) T-47D. Effect of siRNA-mediated BUB1 depletion on radiosensitization was measured in these cell lines. Transient BUB1 siRNA transfection led to moderate radiosensitization with rER 1.0 to 1.2 in (F) SUM159, and (G) MDA-MB-231; After silencing of BUB1, BUB1-WT re-expression rescues the radiosensitization phenotype while BUB1-KD does not in (H) SUM159 and (I) MDA-MB-231. Genomic depletion of BUB1 by CRISPR/Cas9 leads to radiosensitization in (J) SUM159 and (K) MDA-MB-231 cells; Re-expression of BUB1-WT rescues the radiosensitization phenotype in BUB1 CRISPR KO (L) SUM159 and (M) MDA-MB-231 cells but BUB1-KD does not in. P-values were defined as * P≤0.05, ** P≤0.01, and *** P≤0.001.

Effect of siRNA-mediated BUB1 depletion on radiosensitization was measured in all the selected breast cancer cell lines (**Fig. 3F-G; Supplementary Fig. S5**). We observed moderate radiosensitization (rER 1.0 to 1.2) when BUB1 was transiently depleted by siRNA. BUB1 depletion led to a significant reduction in the surviving fraction at 2 Gy (SF-2 Gy). Western blot analyses of total cell lysates following transfection of siRNA revealed that BUB1 could be efficiently repressed. In order to confirm that these effects are mediated by BUB1, we performed the same experiments in SUM159 and MDA-MB-231 (BUB1 depleted) cells with reintroduction of wild-type or kinase dead BUB1 (BUB1-wt, BUB1-kd) (**Fig. 3H-I**). Addition of BUB1-wt restored radioresistance in both the cell lines (rER 0.9) while BUB1-kd addition did not (rER 1.0 to 1.1) and this response was correlated with immunoblotting analyses.

Since transient BUB1 depletion by siRNA did not lead to significant radiation sensitization, we generated BUB1 knockout (BUB1 KO) SUM159 and MDA-MB-231 cell lines by CRISPR-CAS9 RNP transfection. Multiple BUB1 CRISPR clones were validated by Western blotting and Sanger sequencing to confirm complete BUB1 KO (**Supplementary Fig. S6**). Two different BUB1 KO clones for each cell line were used for subsequent experiments. SUM159 BUB1 KO clones demonstrated significant radiation sensitization (clone #18 rER 1.24, clone #48 rER 1.27) (**Fig. 3J**). There was also significant decrease in surviving fractions at 2 Gy (SF-2 Gy) in these clones. Similarly, significant radiation sensitization was observed in MDA-MB-231 BUB1 KO clones (clone #12 rER 1.57, clone #15 rER 1.37) and also significant reduction in surviving fractions at 2 Gy (**Fig. 3K**). To further confirm a role for BUB1 in radiation sensitization, BUB1-wt and BUB1-kd plasmids were transfected in one BUB1 CRISPR KO SUM159 and MDA-MB-231 clone each and clonogenic survival assay was performed (**Fig. 3L-M**). In both the cases, we observed significant radiation sensitization which was reversed when BUB1-wt was expressed (rER 0.9) but not in BUB1-kd expressed cells (rER 1.0), as demonstrated by immunoblotting. The rER value of SUM159 and MDA-MB-231 BUB1 KO clones presented in **Fig. 3L-M** is lower than that of **Fig. 3J-K** due to the toxicity that is commonly observed with the Lipofectamine 2000 transfection reagent, which we used to transfect the BUB1-wt and BUB1-kd plasmids.

### BUB1 inhibition radiosensitizes SUM159 tumor xenografts and prolongs animal survival

To determine the effects of BUB1 inhibition on radiosensitization *in vivo*, xenograft tumors (N= >9-10/arm) were generated by injecting SUM159 cells into the 4^th^ mammary fat pads of female CB17/SCID mice. Mice were randomized to different treatment groups once the tumors reached ∼80 mm^3^. Mice received either BUB1 inhibitor BAY1816032 (25 mg/kg, twice daily for 4 weeks, weekdays only), RT (5Gy X3, 2 days apart), combination or sham irradiation/vehicle (**Fig. 4A**). We initially tested the doses/fractions (2.5 GyX8, 5 GyX3 and 10 GyX1) that yielded similar equivalent dose (EQD2; 21.7-23.3Gy) and biologically effective dose (BED; 32.5-35Gy) using an alpha/beta ratio of 4 which enabled us to explore whether high dose/fraction was more effective than standard fractionation. We observed insignificant benefits of adding 2.5GyX8 and 10 GyX1 radiation with BUB1i while 5GyX3 schema demonstrated superior tumor control (**Supplementary Fig. S7A**) which was selected for the subsequent repeat experiments. In combination treatment arm, RT started 24 h after the first treatment with BUB1i. BAY1816032 with RT significantly reduced tumor growth (**Fig. 4B**) compared with inhibitor or RT alone and significantly extended animal survival (**Fig. 4C-E**). There was no toxicity of BUB1 inhibitor since body weight of experimental animals remained constant during the study period (**Supplementary Fig. S7B**). Immunohistochemical staining of Ki67 (marker for proliferation) from tumors collected at the study end point revealed a significant reduction in Ki67 positivity in combination treatment than either treatment alone which also correlated with H&E staining pattern (**Fig. 4F-G**).

**Fig. 4.**
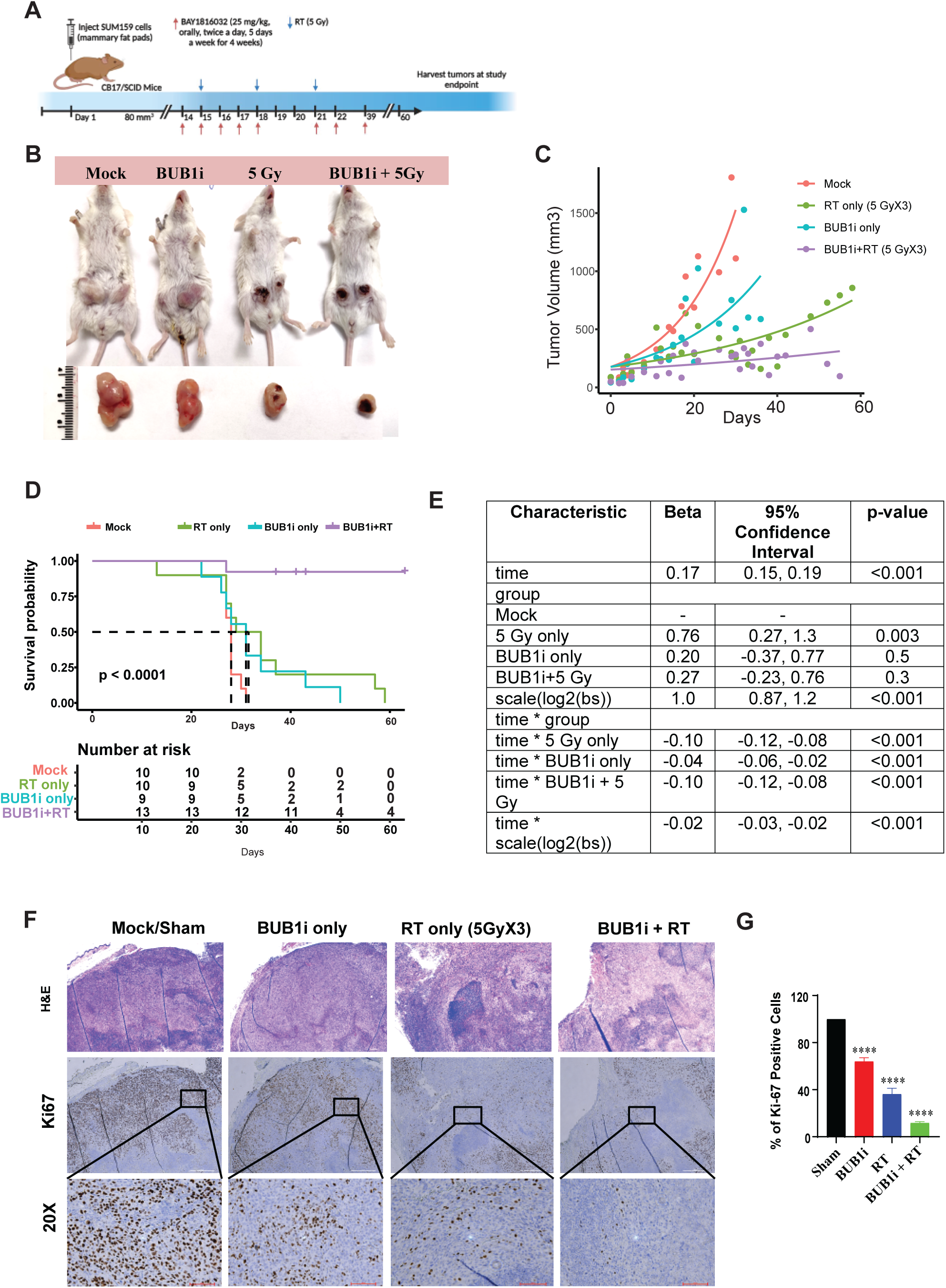
BUB1 inhibition sensitizes SUM159 tumor xenografts to radiation (A) Timeline of the experiment; (B) Representative images of tumor growth in different treatment groups; (C) Combination treatment of BAY1816032 + RT reduces tumor volume in vivo; (D, E) Combination treatment increases tumor volume doubling time in Fox Chase SCID mice; Representative images of (F) H&E staining showing structural changes and Ki67 staining (a proliferation marker) revealed a significant reduction in combination treatment of SUM159 xenografts; (G) Ki-67 plot showing decrease in % of positive cells in combination treatment of BUB1i + RT. P-value was defined as **** P≤0.0001.

Additionally, we generated mammary fat pad tumor xenografts in CB17/SCID mice (N = 4-10/arm) using SUM159 BUB1 CRISPR KO cell line (clone #48). Animals were randomly divided into treatment groups once the tumors established (∼80 mm^3^) and treated with RT (5Gy X3) or sham irradiated (**Fig. 5A**). There was a significant increase in mouse survival in combination treatment group as compared to sham irradiation (**Fig. 5B-E**).

**Fig. 5.**
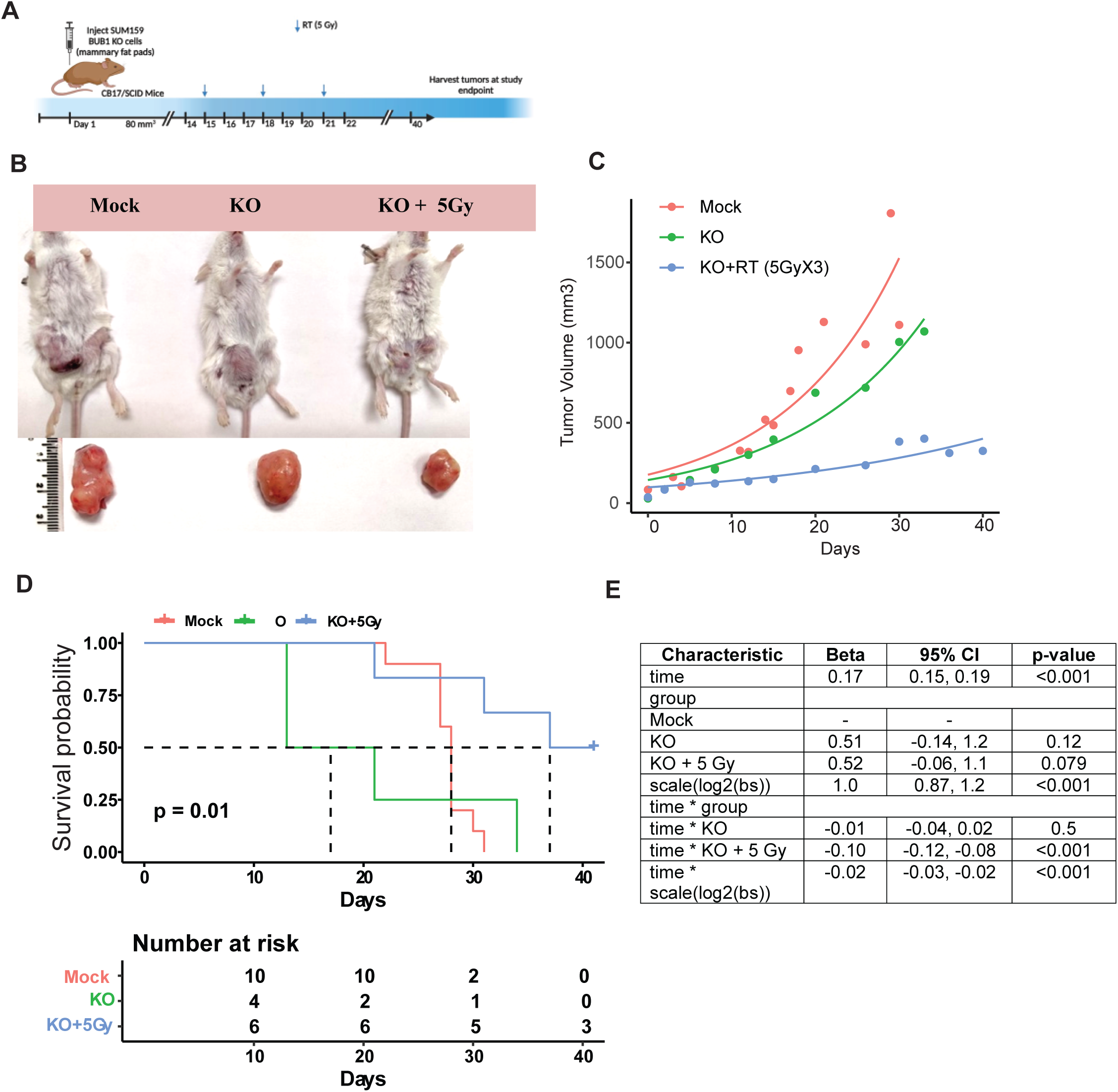
Tumor xenograft of BUB1 CRISPR KO SUM159 cells are sensitive to irradiation. (A) Timeline of the experiment; (B) Representative images of tumor growth in different treatment groups; (C) Treatment of BUB1 KO + RT reduces tumor volume in vivo; (D, E) Treatment of BUB1 KO + RT increases tumor volume doubling time in Fox Chase SCID mice. P-value was defined as * P≤0.05.

### BUB1 inhibition reduces radiation induced DSB repair as visualized by **γ**H2AX foci

We next investigated the effect of BAY1816032 on dsDNA break repair. γH2AX foci (> 10 foci per cell), a marker for unresolved double strand DNA damage was assessed in cells treated with DMSO and 1 μM BAY1816032, either with or without RT (4 Gy) at different time points (30 min, 4 h, 16 h, 24 h). NU7441 (DNAPK inhibitor) was used as a positive control. Representative images are shown of γH2AX (16 h) in SUM159 and MDA-MB-231 cell lines (**Fig. 6A and C**). Non-irradiated cells had fewer γH2AX positive cells. RT induced the formation of γH2AX foci in approximately 40% of cells within 30 mins post-irradiation, peaked at 4 h, gradually decreased by 16 h and reached near baseline levels by 24 h. However, pretreatment with BAY1816032 resulted in a slight increase in the number of foci (approximately 90% of cells) at 30 min post-irradiation and the expression of γH2AX foci continued to remain elevated thereafter; even at 16 and 24 h with a significantly higher number of foci in the BAY1816032 pre-treated group compared to RT alone group (**Fig. 6B and D**). Cells treated with RT alone efficiently repaired the RT-induced dsDNA damage than the combination over the time, suggesting that BUB1 inhibition delayed the RT induced dsDNA break repair efficiency.

**Fig. 6.**
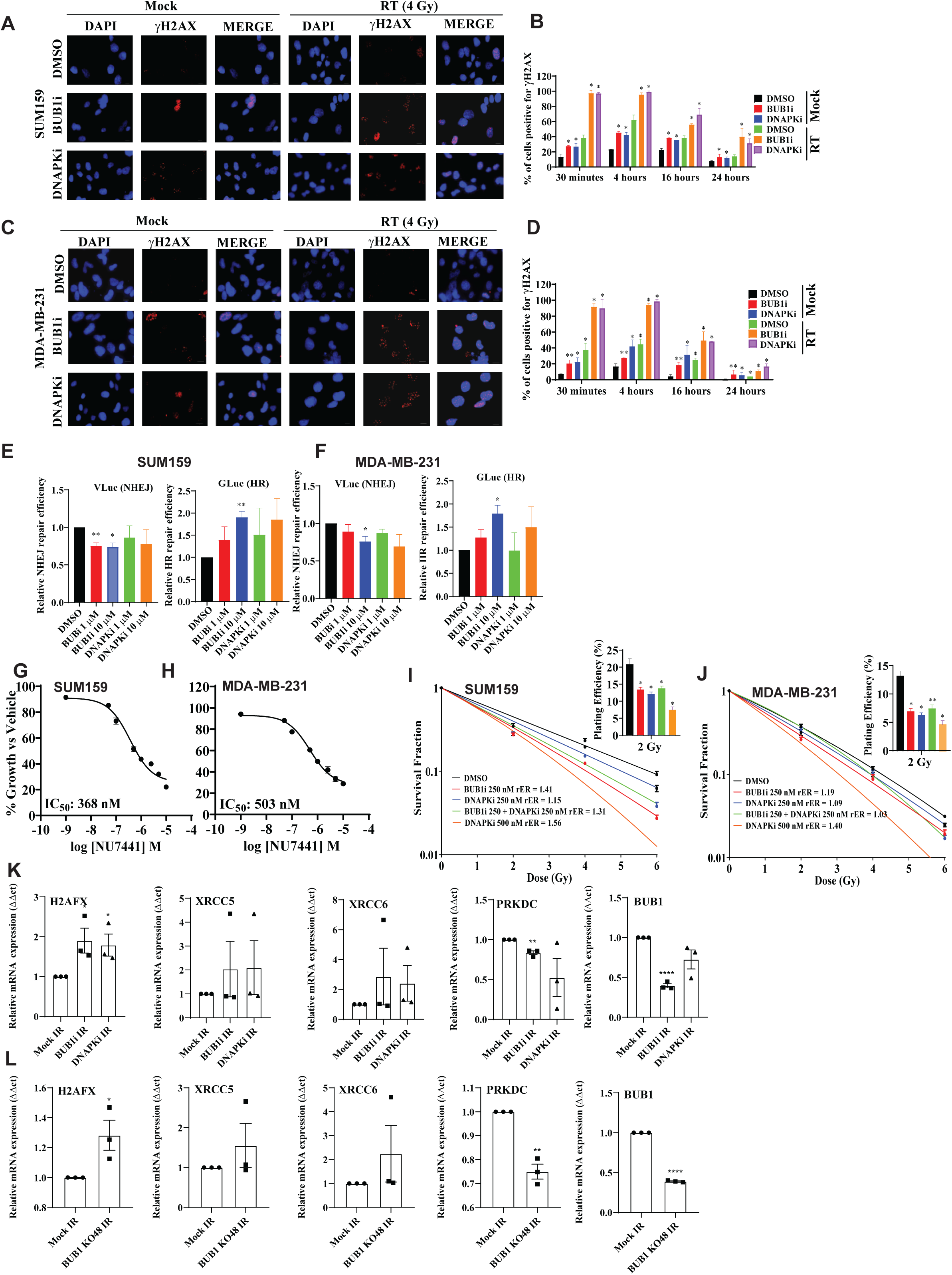
BUB1 ablation radiosensitize through NHEJ. Representative images of (A) SUM159 and (C) MDA-MB-231 γH2AX foci at 16 h. Original magnification, ×63; Combination treatment of BUB1i and RT leads to delayed resolution of γH2AX foci in (B) SUM159, and (D) MDA-MB-231 cell lines. Inhibition of BUB1 kinase function by BAY1816032, at 1 μM and 10 μM, decreases NHEJ efficiency (V Luc) and increases HR efficiency (G Luc) in (E) SUM159, and (F) MDA-MB-231. Effect of DNAPK inhibitor (NU7441) on cell proliferation in TNBC cell lines. NU7441 is cytotoxic to cells at low nanomolar range (G) SUM159, IC_50_: 368 nM; (H) MDA-MB-231, IC_50_: 503 nM; Combination of BAY1816032 and NU7441 does not increase DNAPKcs-mediated radiosensitization in (I) SUM159 (J) MDA-MB-231 cell lines. Inhibition of BUB1 increased transcription of DNA damage genes after radiation. Significant upregulation of *H2AFX* and downregulation of *PRKDC* levels in (K) SUM159 and (L) SUM159 BUB1 CRISPR KO cells were observed. P-values were defined as * P≤0.05, ** P≤0.01, *** P≤0.001, **** P≤0.0001.

We also assessed the dsDNA break repair using BUB1 siRNA in SUM159 and MDA-MB-231 cell lines. Representative images of γH2AX (16 h) are shown in **Supplementary Fig. S8**. Almost all cells treated with BUB1 siRNA were γH2AX foci positive in presence or absence of RT (4 Gy) at 30 min and 4 h. The largest differences were seen at subsequent time points (16 and 24 h) in which BUB1 depletion resulted in persistence of γH2AX foci, whereas the foci began to resolve in presence of BUB1. These results indicate that inhibition of BUB1 kinase activity most likely results in a slower rate of DNA damage repair.

### BUB1 inhibition reduces non-homologous end joining (NHEJ) repair

The two major pathways for repair of DNA DSBs include HR and NHEJ (31). Though either may be involved in repairing dsDNA breaks, earlier reports suggested a potential link between BUB1 expression and NHEJ pathway (10). Thus, we hypothesized that reduced NHEJ repair efficiency is partly responsible for BUB1-mediated radiosensitization and prolonged unresolved dsDNA breaks. Following the induction of a DSB, we used BLRR approach to simultaneously monitor the NHEJ and HR dynamics (23). We aimed to confirm if BUB1 inhibition impacted NHEJ or HR since it has been previously demonstrated that knockdown of BUB1 reduces NHEJ efficiency (10). BLRR transfected cells treated with BAY1816032 at two different concentrations (1 and 10 μM) in presence of RT (4 Gy) led to a decrease in NHEJ (VLuc activity) signal significantly decreased as the GLuc signal (HR activity) increased reciprocally in a dose-dependent manner (**Fig. 6E-F**). NU7441 was used as a positive control. These results indicate that BUB1 inhibition decreases NHEJ-mediated DNA damage repair efficiency and BUB1-mediated radiosensitization may take place through the NHEJ pathway.

### BUB1 inhibition does not increase DNAPKi-mediated radiosensitization

The above results encouraged us to further assess the effect of BUB1 inhibition in combination with a DNAPK specific inhibitor NU7441, which is well-known to impair NHEJ-mediated radiation-induced DSB repair (32). Initially, we investigated the cytotoxicity of NU7441 in SUM159 and MDA-MB-231 cells at 72 h. The IC_50_ value of NU7441 on these cells ranges from 300 – 500 nM (**Fig. 6G-H**). The radiosensitization effects following treatment with a combination of BAY1816032 and NU7441 (250 nM each) in presence of radiation (0, 2, 4, 6 Gy) were assessed using clonogenic survival assays. When combined, BUB1 inhibition does not increase DNAPK inhibitor driven radiosensitization (combination rER ranges from 1 to 1.3) which further confirms that BUB1-mediated radiosensitization takes place through NHEJ pathway (**Fig. 6I-J**). Furthermore, the surviving fraction at 2 Gy (SF-2 Gy) was significantly reduced by the combined effect of BAY1816032 and NU7441 (**Fig. 6I-J; inset plots**) but it was not significantly different than either agent alone.

### Pharmacological and genomic ablation of BUB1 causes increased transcription of DNA damage genes after radiation

Cells were pre-treated with BUB1i, irradiated (4 Gy) 1h after BUB1i and harvested 72h post RT to examine the impact of BUB1 inhibition on NHEJ pathway associated genes by qPCR. The expression of *H2AFX, XRCC5, XRCC6, PRKDC,* and *BUB1* was measured and normalized against *GAPDH*. These results demonstrated an increase in H2AFX, XRCC5, and XRCC6 in BUB1i treated SUM159 (**Fig. 6K**) and MDA-MB-231 cells (**Supplementary Fig. S9, top panel**). We observed significant downregulation of *PRKDC* and BUB1 in BUB1i treated cells. (**Fig. 6K** and **Supplementary Fig. S9**). Similar results were obtained in both BUB1 CRISPR KO cell lines (SUM159 KO #48; **Fig. 6L** and MDA-MB-231 KO #12; **Supplementary S9, bottom panel**) further supporting a role for BUB1 in regulating mRNA levels of key NHEJ genes in response to radiation.

### BUB1 ablation increases DNAPKcs phosphorylation and stabilizes it after irradiation

DNAPK catalytic subunit (DNAPKcs) is a well-known mediator of DNA DSB repair through the activation of NHEJ (33, 34). DNAPK autophosphorylates at Ser2056 (PQR cluster) and Thr2609 (ABCDE cluster) in response to DSB induction (35–37) which may limit or promote DNA end processing during NHEJ (36, 38). Thus, we evaluated if BUB1 ablation had any effect of DNAPK phosphorylation at Ser2056 (S2056) in MDA-MB-231 (**Fig. 7A**) and MDA-MB-468 (**Supplementary S10A**) cells. As expected, radiation treatment led to an increase in DNAPK phosphorylation (pDNAPKcs) at S2056 which was significantly increased in samples that had been pre-treated with BUB1i (**Fig. 7A**). DNAPK inhibitor NU7441 was used as a positive control in parallel experiments. Not surprisingly, pre-treatment with NU7441 almost completely blocked radiation induced DNAPKcs S2056 phosphorylation in these cells (**Fig. 7A** and **Supplementary S10A**). There were no noticeable changes in the expression of KU70, KU80, or total DNAPKcs. Since radiation induced DNAPKcs autophosphorylation can be observed within minutes (25, 35, 36) and phospho-DNAPKcs levels decrease afterwards (37), we next investigated if BUB1 ablation changes pDNAPKcs dynamics following radiation. Cells were pre-treated with BUB1i for 1hr followed by 4 Gy radiation and collected at various intervals (0, 15, 30, and 120 min). We observed that the pre-treatment with BAY1816032 augmented the expression of pDNAPKcs (S2056), which was noticeable up to 2h while pDNAPKcs started to decrease after 30 minutes in the radiation alone group in MDA-MB-231 while it was noticeable in MDA-MB-468 only at 120 minutes in RT only lanes (**Fig. 7B and Supplementary S10B**). This data indicates that BUB1 ablation increases the amplitude and duration of radiation induced pDNAPKcs within a PQR cluster site.

**Fig. 7.**
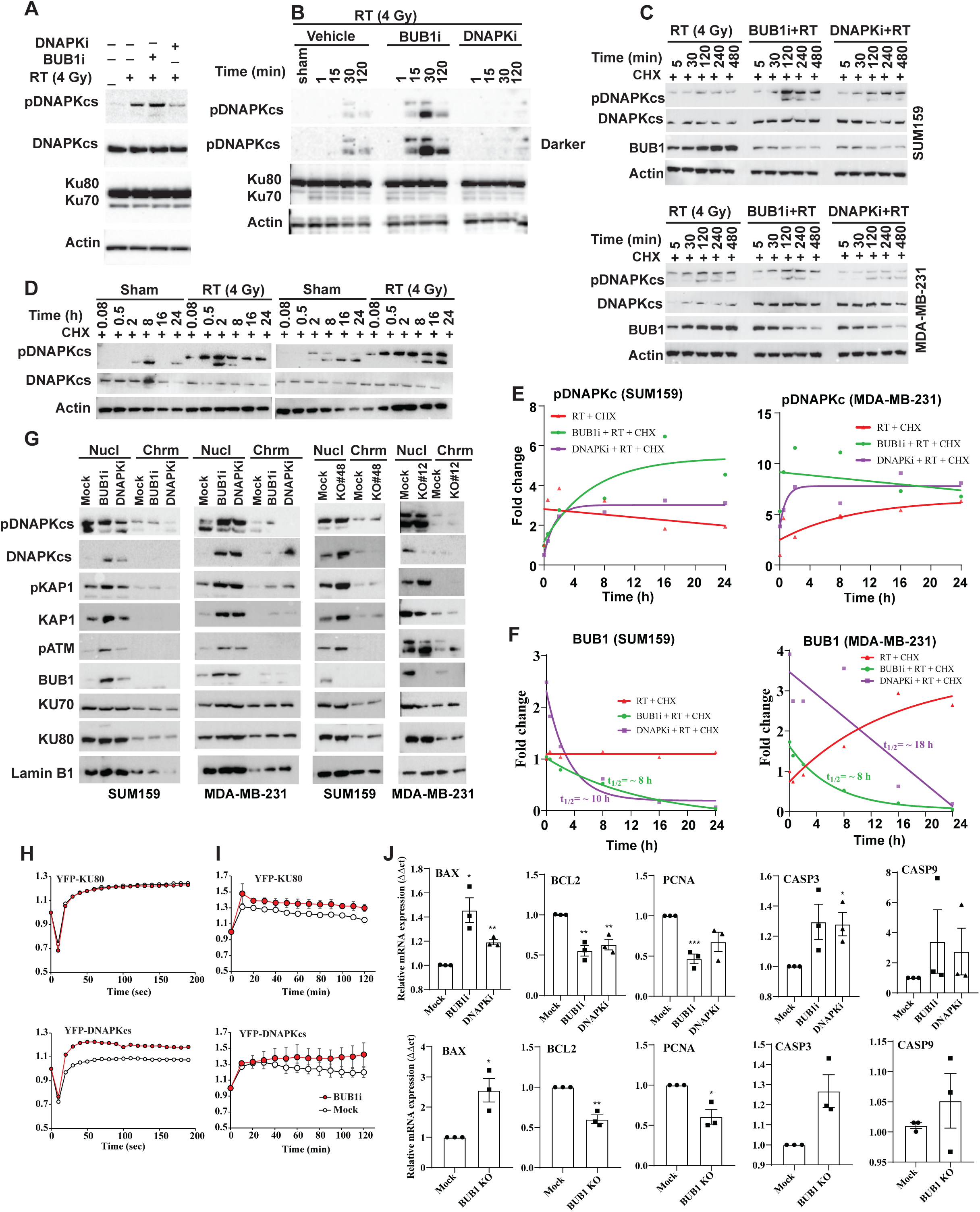
BUB1 ablation leads to increased phosphorylation of DNAPKcs, alters chromatin localization of key NHEJ factors and induces apoptotic cell death upon irradiation. (A) MDA-MB-231 cells were treated with BUB1i or DNAPKi an hour prior to radiation treatment. Cells were harvested 30 minutes post RT (4Gy) and resolved on SDS-PAGE gels and probed with indicated antibodies. (B) MDA-MB-231 cells were treated as (A) and harvested at 1-, 15-, 30– and 120-minutes post-RT and immunoblotted as specified. (C) SUM159 (top panel) MDA-MB-231 cells (bottom panel) were treated with cycloheximide followed by BUB1i or DNAPKi and radiation (4Gy). Total protein lysates were made at the indicated time-points and resolved on gels. (D) BUB1 CRISPR KO SUM159 (left panel) or MDA-MB-231 (right panel) cells were treated with cycloheximide, and radiation and samples were harvested at different time points. (E) Quantitation of pDNAPKcs protein levels in SUM159 and MDA-MB-231 cells (from 7C and other experiments). (F) Quantitation of BUB1 protein levels in SUM159 and MDA-MB-231 cells (from above experiments). (G) Nuclear and chromatin fractions of SUM159 and MDA-MB-231 cells treated with BUB1i, DNAPKi and RT (left) and BUB1 CRISPR KO SUM159 and MDA-MB-231 cells treated with RT (right panels). (H) Effect of BUB1 inhibitor (red circles) on initial recruitment of YFP-tagged KU80 and YFP-DNAPKcs by laser microirradiation in U2OS cells. (I) effect of BUB1 inhibition on the accumulation of YFP-KU80 and YFP-DNAPKcs at laser-induced DSBs for up to 120 minutes. (J) QRT-PCR of BAX, BCL2, PCNA, CASP3 and CASP9 in SUM159 cells treated with BUB1i, DNAPKi and radiation (4 Gy, 72 hours). (I) QRT-PCR of BAX, BCL2, PCNA, CASP3 and CASP9 in SUM159 BUB1 CRISPR cells 72 hours post-irradiation (4 Gy). P-values were defined as * P≤0.05, ** P≤0.01, and *** P≤0.001.

To validate if observed increase in amplitude and duration of pDNAPKcs was due to the stabilization of pDNAPKcs-S2056, we carried out an experiment wherein nascent protein synthesis was blocked by cycloheximide (CHX). MDA-MB-231 and SUM159 cells were treated with CHX, followed by BUB1i, DNAPKi, vehicle/mock and radiation (4Gy). Protein samples were collected at various time points (0 min, 30 min, 2 h, 8 h, 16 h, and 24 h) and resolved on SDS-PAGE gels (**Fig. 7C**). Densitometric analysis yielded half-life (t_1/2_) of pDNAPKcs at >24h in radiation treated samples which significantly increased upon DNAPKi treatment in SUM159 cells. Surprisingly, combination of BUB1i with RT significantly stabilized pDNAPKcs up to the longest time point evaluated (24h) such that t_1/2_ could not be estimated (**Fig. 7C**). These results demonstrate that BUB1 ablation stabilizes radiation induced pDNAPKcs. In BUB1 CRISPR KO cell lines (SUM159 KO#48 and MDA-MB-231 KO#12), DNAPKcs phosphorylation was detectable up to 24 h in the presence of RT further confirming a role for BUB1 in stabilizing pDNAPKcs (i.e., active DNAPKcs) in response to radiation (**Fig. 7D**). Interestingly, we observed that BUB1 protein was stabilized upon radiation treatment (t_1/2_ = ∞, **Fig. 7C and 7E**) which was reversed in cells pre-treated with BAY1816032 (t_1/2_ = 8h, **Fig. 7F**). BUB1 inhibitor at clonogenic concentrations did not affect BUB1 protein levels in MCF10A cells (**Supplementary Fig. S11**).

### BUB1 ablation alters chromatin localization of NHEJ proteins

Chromatin remodeling increases the accessibility of the region surrounding a DNA lesion for proteins involved in DNA damage response and repair(39). DNA damage sensors and early signal transducers are rapidly attracted to damaged DNA sites right after the radiation exposure(40). We postulated that the initial local chromatin relaxation brought about by BUB1 kinase activity is necessary for the rapid loading of the NHEJ machinery to DSBs. To examine this, nuclear and chromatin fractions were isolated 10 min post-DNA damage with 8 Gy RT in BUB1i-treated and BUB1 CRISPR KO SUM159 and MDA-MB-231 cell lines (**Fig. 7G**). We observed an increased recruitment of phospho-DNAPKcs, total-DNAPKcs, and KAP1 in both nuclear and chromatin-enriched fractions suggesting that BUB1 plays a crucial role in the activation and recruitment of key NHEJ proteins to DSBs. There was no change in the enrichment of KU70 and KU80 proteins in these fractions. The hypothesis that BUB1 is necessary for the quick recruitment of the NHEJ factors to DNA damage sites was supported by laser micro-irradiation experimental findings. Since BUB1 interacts with DNAPKcs just after DSB induction (10), we additionally looked at whether BUB1 regulates DNAPKcs at DSBs. Inhibition of BUB1 does not affect the initial recruitment of YFP-KU80 as viewed in **Fig. 7H** (top, 200 sec) while BUB1i results in rapid recruitment of YFP-DNAPKcs to DSBs compared to the vehicle treated cells (**Fig. 7H**, bottom). In contrast, BUB1 inhibition resulted in prolonged retention of KU80 and DNAPKcs at DSBs for up to 120 minutes (**Fig. 7I**). Gene correlation analysis on the METABRIC dataset identified very strong correlation between BUB1 and H2AX (spearman correlation 0.58), PRKDC (0.39), and moderate correlation with XRCC5 (0.05) and XRCC6 (0.29) further corroborating a strong link between BUB1 and NHEJ mediators (**Supplementary S12**).

### BUB1 ablation increases transcription of apoptotic genes after irradiation

Since BUB1 increased radiation induced cell death (**Fig. 2H-M**, and **Fig. 3F-M**) and led to increased loading of key NHEJ factors chromatin fractions (i.e., DNA damage; **Fig. 7G**), we next sought out to elucidate cell death mechanisms mediated by the combination treatment. qRT-PCR for pro-apoptotic, anti-apoptotic and proliferation genes demonstrated significant upregulation of *BAX*, *CASP3* and *CASP9* while significant downregulation of *PCNA* and *BCL2* was observed after BUB1 ablation in SUM159 and MDA-MB-231 cell lines (**Fig. 7J and Supplementary S13**). Gene correlation studies using the METABRIC dataset identified very strong correlation between BUB1 and MKI67 (spearman correlation 0.71), CASP3 (0.44), BAX (0.27), BCL2 (–0.42) and PCNA (0.48) all with p <*** (**Supplementary S13**) further supporting a role for BUB1 in facilitating radiation induced apoptosis.

### BUB1 is overexpressed in tumors and its expression correlates with tumor grade

We examined the expression of BUB1 in breast tumors (N = 202) and compared with normal breast tissues (N = 15). Expression levels of BUB1 protein were graded based on staining intensity and percentage of cells positively stained for BUB1. Levels of immunopositivity were scored as follows: 0 (No staining); 1+ (Weak staining); 2+ (Moderate staining); 3+ (Strong staining). Scores of 0 designated as negative, and scores of 1, 2, and 3 were designated as positive. Examples of BUB1 staining are illustrated in **Fig. 8A** under 4x and 20x magnifications. Immunohistochemical analysis revealed a significantly high BUB1 protein expression in breast tumors compared to normal breast tissue. We observed significant correlation between BUB1 protein expression (staining intensity) and tumor grades and stages (**Fig. 8B**). Furthermore, BUB1 was overexpressed in TNBC (N = 50; *P<0.05*), ER+/PR+ (N = 63; *P<0.001*), ER+/PR+/HER2+ (N = 19; *P<0.05*), ER+ (N = 37; *P<0.05*), ER+/HER2+ (N = 12; *P<0.01*), HER2+ (N = 18; *P<0.0001*), and PR+ (N = 3; *P<0.0001*) compared to normal breast (N = 15). Although, we observed highest BUB1 staining intensity in PR+ tumors, the number of PR+ samples in the current TMA are too small to statistically support the findings.

**Fig. 8.**
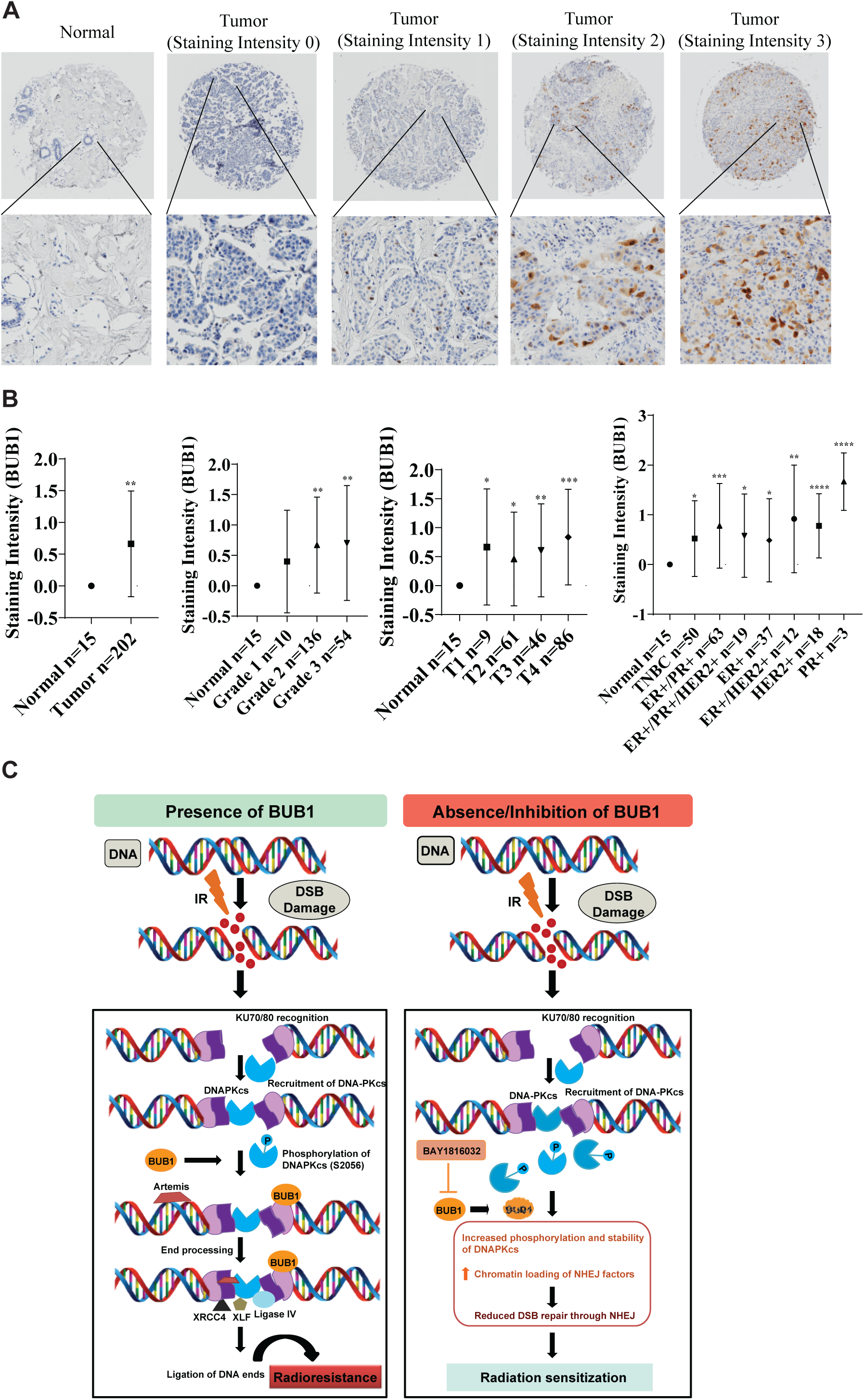
BUB1 is overexpressed in breast tumors. (A) Representative images of BUB1 staining intensity at 4x and 20x magnifications in breast TMA. (B) Quantification of BUB1 staining in breast tumor TMA. (C) Proposed model for a role of BUB1 in mediating radiation induced NHEJ signaling. We propose that radiation induced DNA DSB are repaired efficiently when BUB1 is present (left panel) leading to radiation resistance. In the absence of BUB1 activity or availability, radiation induces hyper phosphorylation of DNAPKcs (Ser2056) and increased binding of NHEJ mediators at the DNA DSB sites (right panel). These NHEJ mediators may not stay on the extended chromatin thus hamper end processing causing radiation-sensitization. P-values were defined as * P≤0.05, ** P≤0.01, *** P≤0.001, and **** P≤0.0001.

## Discussion

BUB1 is a serine/threonine kinase required for optimal DNA damage response as there is increasing evidence that DNA damage response elements and spindle assembly checkpoint components crosstalk (11). We identified BUB1 as a key kinase associated to radiosensitivity in a focused human kinome screen (27). Nevertheless, there is no data that links BUB1 to radiation therapy or DNA damage repair in TNBC. Here, we demonstrate that BUB1-specific inhibitor BAY1816032 radiosensitized TNBC models, a subtype of breast cancer known to have limited treatment options with poorest prognosis (41). Previous studies have shown the advantage of radiation therapy in reducing local recurrence rates, and this was validated in a randomized controlled trial in patients with TNBC (42). However, radioresistance is a major cause of treatment failure or locoregional relapse in TNBC. Here, we provide evidence that BUB1 mediates radiation resistance in TNBC through modulating DNA DSB repair.

Our study showed that BUB1 is overexpressed (differential mRNA levels) in breast cancer with the highest expression in TNBC (**Fig. 1D**). However, BUB1 protein expression (**Fig. 2G**) is found to be slightly different when compared to the differential mRNA expression as delayed protein synthesis may result in reduced mRNA/protein correlations (43, 44). Our findings that BUB1i was effective at a log lower concentration in cancer cells (**Fig. 2A-E**) compared to normal breast epithelial MCF10A cell line (**Fig. 2F**) demonstrate the selectivity and potentially minimal toxicity of BUB1i in future translational studies given it was found to be safe in large animal models (45). Our observations that BUB1 ablation sensitizes TNBC cell lines (SUM159, MDA-MB-231, MDA-MB-468, and BT-549) but not Luminal A subtype (T47D) further support a role for BUB1 in mediating radiation-resistance phenotype in TNBC (27). PI3K family kinases including ATM, ATR, and DNAPK phosphorylate Ser139 in H2AX upon DNA damage which is necessary to sustain the stable association of repair factors at DSB sites (46). ATM phosphorylates BUB1 at Ser314 that activates BUB1 resulting in optimal DNA damage response (11). By re-expressing BUB1-WT and BUB1-KD in BUB1 knockout cells, we confirmed that BUB1 activity plays a role in radiation (**Fig. 3**) and DDR responses (**Fig. 6A-D**). Biochemical or genomic BUB1 ablation radiosensitized SUM159 mouse xenograft model (**Fig. 4** and **Fig. 5**) further corroborating a role for BUB1 in mediating radiation response. Prolonged presence of γH2AX foci after irradiation in BUB1 ablated cells supports earlier reports on delayed or unrepaired DSB after BUB1 ablation (10, 47). The BLRR assay confirmed that BUB1i radiosensitizes TNBC through NHEJ (**Fig. 6E-F**) which was further confirmed by no increase in DNAPKi mediated radiosensitization by BUB1i (**Fig. 6I-J**). Similar observations with DNAPKi have been reported (48).

Recent evidence has shown that the DNAPKcs and DNA methyltransferase inhibitors are effective at sensitizing TNBC to PARPi and radiation (49). Autophosphorylation of DNAPKcs at Ser2056, a known autophosphorylation site within the PQR cluster regulates DNA end processing and possibly DSB repair pathway choice (50, 51). Surprisingly, we identified that BUB1 ablation increased the level and amplitude of pDNAPKcs-S2056 following radiation. The pDNAPKcs was stabilized till the longest time point evaluated (24hr). Our observations support previous studies which demonstrated higher or persistent pDNAPKcs following DNAPKi, ATMi with IR or other DNA damaging agents (35, 51–53). Although, we have not confirmed the mechanism of pDNAPKcs stabilization by BUB1, we are tempted to speculate that known (RNF144A (54), MARCH5 (55), CRL4ADTL (56)) or yet unrelated E3-ubiquitin ligase(s) may be involved. Although tumor suppressor protein P53 (p53) has been linked to BUB1 expression (57), the data has been lacking that demonstrated BUB1 protein regulation by radiotherapy. Our cycloheximide chase assays clearly demonstrates that BUB1 is stabilized upon RT while pretreatment with BUB1i or DNAPKi reverses this and causes BUB1 degradation after irradiation (**Fig. 7C, 7F**). Future studies will determine the mechanism of BUB1 regulation by radiation.

Chromatin remodeling increases the accessibility of DNA damage response and repair proteins in area around a DNA lesion (39). Phosphorylation of H2AX and KAP1 are key steps that enhance chromatin relaxation and allow the recruitment of the DDR machinery to a DSB (58, 59). Our findings (**Fig. 7G**) that BUB1 ablation causes increased loading of pDNAPKc, pKAP1, KAP1, and pATM to chromatin fractions and alters the recruitment of YFP-KU80 and YFP-DNAPKcs (**Fig. 7H-I**) support a role for BUB1 in this step. Lu et al., identified a role for DNAPK kinase activity wherein attenuated chromatin recruitment of MRN complex was detected in DNAPK-KD or null (-/-) cells (25). Additionally, they observed decreased localization of NHEJ factors including LIG4, XRCC4 and XLF in chromatin fractions in these cells further supporting a role for DNAPKcs activity in NHEJ. The above data supports our hypothesis wherein BUB1 mediate radiation induced NHEJ through regulating activation (phosphorylation) of DNAPKcs thus chromatin relaxation and access of NHEJ factors. Taken together, our results provide evidence to the hypothesis that BUB1 is necessary for the quick recruitment of the NHEJ factors to DNA damage sites (**Fig. 7G-I**). In future, we will perform in-depth mechanistic studies such as micronuclease digestion, immunoprecipitation (CO-IP), and proximality ligation assay (PLA) to confirm a role for BUB1 in this step of NHEJ. Since mutagenesis of DNAPKcs Ser2056 confirmed that it limits end-processing (51), additionally we will evaluate if BUB1 ablation impacts this process. Because DNAPKcs inhibition or depletion leads to reduction in NHEJ and reciprocally a shift to HR (60, 61) and since certain phosphorylation in DNAPKcs promote HR while inhibiting NHEJ (60), it would be interesting to see if these phospho sites in DNAPKcs are affected by BUB1 and thus downstream signaling (HR). Moreover, ATM phosphorylates members of MRN complex (that initiates HR cascade) (62), NHEJ factors (including DNAPKcs(63), and H2AX) and is shown to phosphorylate BUB1 at Ser314 in response to irradiation (11), it would be fascinating to investigate if BUB1 indeed affects HR response through ATM or some other mechanism.

BUB1 transcripts are significantly higher in breast cancer cell lines and in high-grade primary breast cancer tissues compared to normal mammary epithelial cells, or in normal breast tissues (64). High BUB1 expression (transcript) correlates with extremely poor outcome in breast cancer (65, 66). Our meta-analysis that BUB1 expression significantly correlates with Ki67 (**Supplementary Fig. S12**) supports earlier findings (65–67) and signifies our in-vivo observations that tumors harvested from mice treated with a combination of BUB1i and radiation have statistically significant reduction in Ki67 (**Fig. 4F-G**) or PCNA in cells (**Fig. 7J**). Our TMA analysis found strong correlation between BUB1 protein expression and tumor grade (**Fig. 8A-B**) and identified high BUB1 expression in TNBC samples. Our BUB1 immunostaining TMA data support earlier findings wherein nuclear BUB1 staining was found to strongly correlate with stage, pathological tumor factors, lymph node metastasis, distant metastasis, histological grade, and proliferation (67). In future it will be important to assess if BUB1 protein expression correlates with treatment naïve or radioresistant-recurrence cases. Based on our data that BUB1 is stabilized upon radiation treatment (**Fig. 7C**) we speculate higher BUB1 expression in radiation resistant, recurrent cases compared to treatment naïve cases.

Our results are consistent with previous reports where knockdown of BUB1 was demonstrated to prolong γH2AX foci, comet tail as well as hypersensitivity in response to ionizing radiation (11). Since BUB1 co-localizes with 53BP1 (10) and interacts with NHEJ factors (10) and we identified that BUB1 ablation increases the amplitude and duration of DNAPKcs phosphorylation and increases chromatin localization of key NHEJ factors, we describe a model on BUB1’s role in NHEJ (Graphical Abstract, **Fig. 8C**). In the presence of BUB1 (left panel), the NHEJ is efficient and can repair radiation induced DSB thus causes radio-resistance. On the other hand, BUB1 inhibition or depletion causes increased phosphorylation of DNAPKcs and increased binding of NHEJ factors at the DSB sites (right panel). These NHEJ mediators do not stay on the extended/open chromatin required for proper end processing and ligation of the DNA ends. This leads to reduced NHEJ repair leading to radiation-sensitization. DNAPKcs phosphorylation is essential for its dissociation from Ku bound DNA (42, 68, 69). Although we observed increased binding of NHEJ factors at the chromatin following BUB1i+RT (10 minutes post RT), we cannot rule out that these factors fall off at a later time without repairing broken DNA ends (limited end processing) as has been demonstrated using DNAPKcs phospho-site mutants (69). Taken together, our data demonstrate that BUB1 is overexpressed in breast cancer including TNBC and BUB1 ablation leads to radiosensitization through regulating DNAPKcs phosphorylation and chromatin localization of key NHEJ factors. Our findings strongly support nomination of BUB1 as a potential biomarker and a therapeutic target for radiosensitization in TNBC.

## Declarations

### Ethics approval and consent to participate

Experimental animals were housed and handled in accordance with protocols approved by IACUC of Henry Ford Health (protocol # 00001298).

### Consent for publication

Not applicable.

### Availability of Data and Materials

Data available from the corresponding author on reasonable request.

## Author contributions

SN conceived and designed the study; SS and ST performed the experiments; WMC performed laser microirradiation experiments, CS performed the bioinformatic analysis; SN, SS, and ST wrote the original manuscript; OH analyzed the immunohistological staining of BUB1 on TMA slides; SS, ST, OH, FS, SLB, BM, MDG, CS, EW, WMC, AD and SN interpreted the data; SS, ST, OH, FS, SLB, BM, MDG, CS, EW, WMC, AD and SN revised the manuscript. All authors approved the final manuscript.

## List of Abbreviations

BC: Breast cancer
BLRR: Bioluminescent repair reporter
BUB1: Budding uninhibited by benzimidazoles-1
CAM: Chick chorioallantoic membrane
CHX: Cycloheximide
DMEM: Dulbecco’s modified eagle medium
DMSO: Dimethyl sulfoxide
DNAPK: DNA dependent protein kinase
DSBs: Double-strand breaks
ECL: Electrogenerated chemiluminescence
EDTA: Ethylenediaminetetraacetic acid
ER: Estrogen receptor
FBS: Fetal bovine serum
GEO: Gene expression omnibus
HER2: Human epidermal growth factor receptor 2
HR: Homologous recombination
IACUC: Institutional Animal Care and Use Committee
IDT: Integrated DNA technologies
KAP1: KRAB-associated protein 1
KD: Kinase-dead
KO: knockout
LMM: Linear mixed models
MRN: Mre11, Rad50 and Nbs1
NHEJ: Non-homologous end joining
PARPi: Poly (ADP-ribose) polymerase inhibitor
PE: Plating efficiency
PEG: Polyethylene glycol
PLA: Proximality ligation assay
POLβ: DNA polymerase-beta
PR: Progesterone receptor
PVDF: polyvinylidene difluoride
qPCR: Quantitative polymerase chain reaction
rER: Radiation enhancement ratio
RNP: Ribonucleoprotein
RT: Radiotherapy
SEM: Standard error of the mean
SF: Survival fraction
SSBs: Single-strand breaks
TCGA: The Cancer Genome Atlas
TMA: Tissue microarray
TNBC: Triple-negative breast cancer
WT: Wild-type

## Acknowledgements

The authors thank Grahm Valadie for help with animal treatment using SARRP and Katheryn Meek (MSU) with data interpretation. We thank Transgenics and CRISPR (TGEF) core at MSU for help with BUB1 gRNA design and Histology core-HFH for immunohistological staining. We thank Pin Li and Sunita Ghosh from Public Health Sciences for statistical analyses. Grant support to SN (NIH/NCI R21 CA252010-01A1), Henry Ford Cancer Institute (HFCI) and Henry Ford Health Research Administration Start Up grant, HFH Proposal Development Award, HFH Near Miss Award, HFH Radiation Oncology Start Up grant), SS (HFCI Translational Oncology Postdoctoral Fellowship).

## Additional Materials

Additional File 1: Supplementary Figures and Tables

## Supplementary Figure/Tables Legends

**Table S1**: List of mutated genes in the TNBC cell lines.

**Table S2**: Guide RNA (gRNA) sequences used to knock out BUB1, primer sequences for PCR amplification of BUB1-edited section, and primer sequence for Sanger sequencing.

**Table S3**: List of antibodies used for Western Blotting/Immunohistochemical/Immunofluorescence studies.

**Table S4**: Primer sequences used in quantitative PCR (qRT-PCR) analysis.

**Fig. S5**: Clonogenic assays using BUB1 siRNA and RT in (A) MDA-MB-468, (B) BT-549, (C) T-47D cell lines. PRKDC siRNA is used as a positive control.

**Fig. S6**: (A) CRISPR-CAS9 RNP transfection method was utilized to knock out BUB1 (B) BUB1 knockouts were confirmed through Immunoblotting in SUM159, and MDA-MB-231 cell lines followed by (C) PCR amplification and (D) Sanger Sequencing to further validate the BUB1 KO’s (E) Sanger Sequencing chromatograms of SUM159 BUB1 KO #48 and #18, and MDA-MB-231 KO #12 and #15.

**Fig. S7**: (A) Initial radiation dose-response studies in SUM159 tumor xenograft in CB17 SCID mice. SUM159 xenograft mammary fat pad tumors were conformally irradiated at 2.5 GyX8, 5 Gy X3 or 10 GyX1 by SARRP (light blue curves). Additionally, mice were treated with a BUB1 inhibitor (25 mg/kg, orally, twice daily, 5 days/week for 4 weeks) along with radiation (red curves). (B) A spaghetti plot for animal body weight change during the treatment.

**Fig. S8**: Immunofluorescence studies using BUB1 siRNA and RT (16 h time point) in (a) SUM159 and (b) MDA-MB-231 cell lines.

**Fig. S9**: qRT-PCR of NHEJ pathway related genes in MDA-MB-231 cell line with BUB1i (top panel) and BUB1 CRISPR-KO #12.

**Fig. S10**: Effect of BUB1 inhibition with IR on DNAPKcs phosphorylation using Immunoblotting in (A) MDA-MB-468 cell line, and (B) shown at different time points up to 2 h.

**Fig. S11:** The effect of BUB1 inhibitor (BAY1816032) on BUB1 protein levels in normal mammary epithelial cell line MCF 10A. The cells were treated for 1 hour with the same doses of BUB1i that were used for the colony formation assays (250 nM, 500 nM and 1000 nM).

**Fig. S12**: mRNA expression plots showing correlation of BUB1 vs. NHEJ pathway-related genes, apoptotic, and proliferation genes in Breast cancer (METABRIC, 2509 samples) from cBIOPORTAL.

**Fig. S13**: qRT-PCR of apoptotic and proliferation genes in MDA-MB-231 cell line with (A) BUB1i and (B) BUB1 CRISPR-KO #12

## References

1. Kyndi M, Sorensen FB, Knudsen H, Overgaard M, Nielsen HM, Overgaard J, et al. Estrogen receptor, progesterone receptor, HER-2, and response to postmastectomy radiotherapy in high-risk breast cancer: the Danish Breast Cancer Cooperative Group. J Clin Oncol. 2008;26(9):1419–26.

2. Mladenov E, Magin S, Soni A, Iliakis G. DNA double-strand break repair as determinant of cellular radiosensitivity to killing and target in radiation therapy. Frontiers in oncology. 2013;3:113.

3. Morgan MA, Lawrence TS. Molecular Pathways: Overcoming Radiation Resistance by Targeting DNA Damage Response Pathways. Clin Cancer Res. 2015;21(13):2898–904.

4. Britton S, Coates J, Jackson SP. A new method for high-resolution imaging of Ku foci to decipher mechanisms of DNA double-strand break repair. The Journal of cell biology. 2013;202(3):579–95.

5. Ahnesorg P, Smith P, Jackson SP. XLF interacts with the XRCC4-DNA ligase IV complex to promote DNA nonhomologous end-joining. Cell. 2006;124(2):301–13.

6. Chapman JR, Taylor MR, Boulton SJ. Playing the end game: DNA double-strand break repair pathway choice. Mol Cell. 2012;47(4):497–510.

7. Gupta A, Hunt CR, Chakraborty S, Pandita RK, Yordy J, Ramnarain DB, et al. Role of 53BP1 in the regulation of DNA double-strand break repair pathway choice. Radiat Res. 2014;181(1):1–8.

8. Berry MR, Fan TM. Target-Based Radiosensitization Strategies: Concepts and Companion Animal Model Outlook. Frontiers in oncology. 2021;11:768692.

9. Sriramulu S, Thoidingjam S, Brown SL, Siddiqui F, Movsas B, Nyati S. Molecular targets that sensitize cancer to radiation killing: From the bench to the bedside. Biomed Pharmacother. 2022;158:114126.

10. Jessulat M, Malty RH, Nguyen-Tran DH, Deineko V, Aoki H, Vlasblom J, et al. Spindle Checkpoint Factors Bub1 and Bub2 Promote DNA Double-Strand Break Repair by Nonhomologous End Joining. Mol Cell Biol. 2015;35(14):2448–63.

11. Yang C, Wang H, Xu Y, Brinkman KL, Ishiyama H, Wong ST, et al. The kinetochore protein Bub1 participates in the DNA damage response. DNA Repair (Amst). 2012;11(2):185–91.

12. Komura K, Inamoto T, Tsujino T, Matsui Y, Konuma T, Nishimura K, et al. Increased BUB1B/BUBR1 expression contributes to aberrant DNA repair activity leading to resistance to DNA-damaging agents. Oncogene. 2021;40(43):6210–22.

13. Nyati S, Schinske-Sebolt K, Pitchiaya S, Chekhovskiy K, Chator A, Chaudhry N, et al. The kinase activity of the Ser/Thr kinase BUB1 promotes TGF-beta signaling. Sci Signal. 2015;8(358):ra1.

14. Nyati S, Gregg B, Xu JQ, Young G, Kimmel L, Mukesh N, et al. TGFBR2 mediated phosphorylation of BUB1 at Ser-318 is required for transforming growth factor-beta signaling. Cancer research. 2019;79(13).

15. Nyati S, Gregg BS, Xu J, Young G, Kimmel L, Nyati MK, et al. TGFBR2 mediated phosphorylation of BUB1 at Ser-318 is required for transforming growth factor-beta signaling. Neoplasia (New York, NY. 2020;22(4):163–78.

16. Tang ZY, Shu HJ, Oncel D, Chen S, Yu HT. Phosphorylation of Cdc20 by Bub1 provides a catalytic mechanism for APC/C inhibition by the spindle checkpoint. Mol Cell. 2004;16(3):387–97.

17. Tang ZY, Sun YX, Harley SE, Zou H, Yu HT. Human Bub1 protects centromeric sister-chromatid cohesion through Shugoshin during mitosis. Proceedings of the National Academy of Sciences of the United States of America. 2004;101(52):18012–7.

18. Yu H, Tang Z. Bub1 multitasking in mitosis. Cell cycle (Georgetown, Tex. 2005;4(2):262–5.

19. Neve RM, Chin K, Fridlyand J, Yeh J, Baehner FL, Fevr T, et al. A collection of breast cancer cell lines for the study of functionally distinct cancer subtypes. Cancer cell. 2006;10(6):515–27.

20. Hatzis C, Pusztai L, Valero V, Booser DJ, Esserman L, Lluch A, et al. A genomic predictor of response and survival following taxane-anthracycline chemotherapy for invasive breast cancer. JAMA. 2011;305(18):1873–81.

21. Speers C, Zhao SG, Kothari V, Santola A, Liu M, Wilder-Romans K, et al. Maternal Embryonic Leucine Zipper Kinase (MELK) as a Novel Mediator and Biomarker of Radioresistance in Human Breast Cancer. Clin Cancer Res. 2016;22(23):5864–75.

22. Zhao SG, Shilkrut M, Speers C, Liu M, Wilder-Romans K, Lawrence TS, et al. Development and validation of a novel platform-independent metastasis signature in human breast cancer. PLoS One. 2015;10(5):e0126631.

23. Chien JC, Tabet E, Pinkham K, da Hora CC, Chang JC, Lin S, et al. A multiplexed bioluminescent reporter for sensitive and non-invasive tracking of DNA double strand break repair dynamics in vitro and in vivo. Nucleic acids research. 2020;48(17):e100.

24. So S, Davis AJ, Chen DJ. Autophosphorylation at serine 1981 stabilizes ATM at DNA damage sites. The Journal of cell biology. 2009;187(7):977–90.

25. Lu H, Saha J, Beckmann PJ, Hendrickson EA, Davis AJ. DNA-PKcs promotes chromatin decondensation to facilitate initiation of the DNA damage response. Nucleic acids research. 2019;47(18):9467–79.

26. Lu H, Shamanna RA, de Freitas JK, Okur M, Khadka P, Kulikowicz T, et al. Cell cycle-dependent phosphorylation regulates RECQL4 pathway choice and ubiquitination in DNA double-strand break repair. Nat Commun. 2017;8(1):2039.

27. Speers C, Zhao S, Liu M, Bartelink H, Pierce LJ, Feng FY. Development and Validation of a Novel Radiosensitivity Signature in Human Breast Cancer. Clin Cancer Res. 2015;21(16):3667–77.

28. Servant N, Bollet MA, Halfwerk H, Bleakley K, Kreike B, Jacob L, et al. Search for a gene expression signature of breast cancer local recurrence in young women. Clin Cancer Res. 2012;18(6):1704–15.

29. Goodwin JF, Knudsen KE. Beyond DNA repair: DNA-PK function in cancer. Cancer Discov. 2014;4(10):1126–39.

30. Yue X, Bai C, Xie D, Ma T, Zhou PK. DNA-PKcs: A Multi-Faceted Player in DNA Damage Response. Frontiers in genetics. 2020;11:607428.

31. Mao Z, Bozzella M, Seluanov A, Gorbunova V. DNA repair by nonhomologous end joining and homologous recombination during cell cycle in human cells. Cell cycle (Georgetown, Tex. 2008;7(18):2902–6.

32. Dong J, Ren Y, Zhang T, Wang Z, Ling CC, Li GC, et al. Inactivation of DNA-PK by knockdown DNA-PKcs or NU7441 impairs non-homologous end-joining of radiation-induced double strand break repair. Oncol Rep. 2018;39(3):912–20.

33. Chang HHY, Pannunzio NR, Adachi N, Lieber MR. Non-homologous DNA end joining and alternative pathways to double-strand break repair. Nat Rev Mol Cell Biol. 2017;18(8):495–506.

34. Kurimasa A, Kumano S, Boubnov NV, Story MD, Tung CS, Peterson SR, et al. Requirement for the kinase activity of human DNA-dependent protein kinase catalytic subunit in DNA strand break rejoining. Mol Cell Biol. 1999;19(5):3877–84.

35. Neal JA, Sugiman-Marangos S, VanderVere-Carozza P, Wagner M, Turchi J, Lees-Miller SP, et al. Unraveling the complexities of DNA-dependent protein kinase autophosphorylation. Mol Cell Biol. 2014;34(12):2162–75.

36. Cui X, Yu Y, Gupta S, Cho YM, Lees-Miller SP, Meek K. Autophosphorylation of DNA-dependent protein kinase regulates DNA end processing and may also alter double-strand break repair pathway choice. Mol Cell Biol. 2005;25(24):10842–52.

37. Chan DW, Chen BP, Prithivirajsingh S, Kurimasa A, Story MD, Qin J, et al. Autophosphorylation of the DNA-dependent protein kinase catalytic subunit is required for rejoining of DNA double-strand breaks. Genes Dev. 2002;16(18):2333–8.

38. Block WD, Yu Y, Merkle D, Gifford JL, Ding Q, Meek K, et al. Autophosphorylation-dependent remodeling of the DNA-dependent protein kinase catalytic subunit regulates ligation of DNA ends. Nucleic acids research. 2004;32(14):4351–7.

39. Price BD, D’Andrea AD. Chromatin remodeling at DNA double-strand breaks. Cell. 2013;152(6):1344–54.

40. Harper JW, Elledge SJ. The DNA damage response: ten years after. Mol Cell. 2007;28(5):739–45.

41. Zagami P, Carey LA. Triple negative breast cancer: Pitfalls and progress. NPJ Breast Cancer. 2022;8(1):95.

42. Abdulkarim BS, Cuartero J, Hanson J, Deschenes J, Lesniak D, Sabri S. Increased risk of locoregional recurrence for women with T1-2N0 triple-negative breast cancer treated with modified radical mastectomy without adjuvant radiation therapy compared with breast-conserving therapy. J Clin Oncol. 2011;29(21):2852–8.

43. Gedeon T, Bokes P. Delayed protein synthesis reduces the correlation between mRNA and protein fluctuations. Biophys J. 2012;103(3):377–85.

44. Liu Y, Beyer A, Aebersold R. On the Dependency of Cellular Protein Levels on mRNA Abundance. Cell. 2016;165(3):535–50.

45. Siemeister G, Mengel A, Fernandez-Montalvan AE, Bone W, Schroder J, Zitzmann-Kolbe S, et al. Inhibition of BUB1 Kinase by BAY 1816032 Sensitizes Tumor Cells toward Taxanes, ATR, and PARP Inhibitors In Vitro and In Vivo. Clinical Cancer Research. 2019;25(4):1404–14.

46. Griesbach E, Schlackow M, Marzluff WF, Proudfoot NJ. Dual RNA 3’-end processing of H2A.X messenger RNA maintains DNA damage repair throughout the cell cycle. Nat Commun. 2021;12(1):359.

47. Morales AG, Pezuk JA, Brassesco MS, de Oliveira JC, de Paula Queiroz RG, Machado HR, et al. BUB1 and BUBR1 inhibition decreases proliferation and colony formation, and enhances radiation sensitivity in pediatric glioblastoma cells. Childs Nerv Syst. 2013;29(12):2241–8.

48. Benjamin C, Chandler LM, Cassandra L. Ritter, Meilan Liu, Meleah Cameron, Kari Wilder-Romans, Amanda Zhang, Andrea M. Pesch, Anna R. Michmerhuizen, Nicole Hirsh, Marlie Androsiglio, S. Tanner Ward, Eric Olsen, Yashar S Niknafs, Sofia D. Merajver, Dafydd Thomas, Powel H. Brown, Theodore S Lawrence, Shyam Nyati, Lori J. Pierce, Arul M. Chinnaiyan, Corey Speers. TTK inhibition radiosensitizes basal-like breast cancer through impaired homologous recombination. Journal of Clinical Investigation. 2019;accepted.

49. Fok JHL, Ramos-Montoya A, Vazquez-Chantada M, Wijnhoven PWG, Follia V, James N, et al. AZD7648 is a potent and selective DNA-PK inhibitor that enhances radiation, chemotherapy and olaparib activity. Nat Commun. 2019;10(1):5065.

50. Mohiuddin IS, Kang MH. DNA-PK as an Emerging Therapeutic Target in Cancer. Frontiers in oncology. 2019;9:635.

51. Jiang W, Crowe JL, Liu X, Nakajima S, Wang Y, Li C, et al. Differential phosphorylation of DNA-PKcs regulates the interplay between end-processing and end-ligation during nonhomologous end-joining. Mol Cell. 2015;58(1):172–85.

52. Quanz M, Chassoux D, Berthault N, Agrario C, Sun JS, Dutreix M. Hyperactivation of DNA-PK by double-strand break mimicking molecules disorganizes DNA damage response. PLoS One. 2009;4(7):e6298.

53. Wang Y, Xu H, Liu T, Huang M, Butter PP, Li C, et al. Temporal DNA-PK activation drives genomic instability and therapy resistance in glioma stem cells. JCI Insight. 2018;3(3).

54. Ho SR, Mahanic CS, Lee YJ, Lin WC. RNF144A, an E3 ubiquitin ligase for DNA-PKcs, promotes apoptosis during DNA damage. Proceedings of the National Academy of Sciences of the United States of America. 2014;111(26):E2646–55.

55. Heo J, Park YJ, Kim Y, Lee HS, Kim J, Kwon SH, et al. Mitochondrial E3 ligase MARCH5 is a safeguard against DNA-PKcs-mediated immune signaling in mitochondria-damaged cells. Cell death & disease. 2023;14(12):788.

56. Feng M, Wang Y, Bi L, Zhang P, Wang H, Zhao Z, et al. CRL4A(DTL) degrades DNA-PKcs to modulate NHEJ repair and induce genomic instability and subsequent malignant transformation. Oncogene. 2021;40(11):2096–111.

57. Elango R, Vishnubalaji R, Shaath H, Alajez NM. Transcriptional alterations of protein coding and noncoding RNAs in triple negative breast cancer in response to DNA methyltransferases inhibition. Cancer Cell Int. 2021;21(1):515.

58. Ziv Y, Bielopolski D, Galanty Y, Lukas C, Taya Y, Schultz DC, et al. Chromatin relaxation in response to DNA double-strand breaks is modulated by a novel ATM– and KAP-1 dependent pathway. Nat Cell Biol. 2006;8(8):870–6.

59. Nakamura AJ, Rao VA, Pommier Y, Bonner WM. The complexity of phosphorylated H2AX foci formation and DNA repair assembly at DNA double-strand breaks. Cell cycle (Georgetown, Tex. 2010;9(2):389–97.

60. Neal JA, Dang V, Douglas P, Wold MS, Lees-Miller SP, Meek K. Inhibition of homologous recombination by DNA-dependent protein kinase requires kinase activity, is titratable, and is modulated by autophosphorylation. Mol Cell Biol. 2011;31(8):1719–33.

61. Neal JA, Meek K. Choosing the right path: does DNA-PK help make the decision? Mutat Res. 2011;711(1-2):73–86.

62. Blackford AN, Jackson SP. ATM, ATR, and DNA-PK: The Trinity at the Heart of the DNA Damage Response. Mol Cell. 2017;66(6):801–17.

63. Lu H, Zhang Q, Laverty DJ, Puncheon AC, Augustine MM, Williams GJ, et al. ATM phosphorylates the FATC domain of DNA-PKcs at threonine 4102 to promote non-homologous end joining. Nucleic acids research. 2023;51(13):6770–83.

64. Yuan B, Xu Y, Woo JH, Wang Y, Bae YK, Yoon DS, et al. Increased expression of mitotic checkpoint genes in breast cancer cells with chromosomal instability. Clin Cancer Res. 2006;12(2):405–10.

65. Dai H, van’t Veer L, Lamb J, He YD, Mao M, Fine BM, et al. A cell proliferation signature is a marker of extremely poor outcome in a subpopulation of breast cancer patients. Cancer research. 2005;65(10):4059–66.

66. Chen DL, Cai JH, Wang CCN. Identification of Key Prognostic Genes of Triple Negative Breast Cancer by LASSO-Based Machine Learning and Bioinformatics Analysis. Genes (Basel). 2022;13(5).

67. Takagi K, Miki Y, Shibahara Y, Nakamura Y, Ebata A, Watanabe M, et al. BUB1 immunolocalization in breast carcinoma: its nuclear localization as a potent prognostic factor of the patients. Horm Cancer. 2013;4(2):92–102.

68. Meek K, Douglas P, Cui X, Ding Q, Lees-Miller SP. trans Autophosphorylation at DNA-dependent protein kinase’s two major autophosphorylation site clusters facilitates end processing but not end joining. Mol Cell Biol. 2007;27(10):3881–90.

69. Uematsu N, Weterings E, Yano K, Morotomi-Yano K, Jakob B, Taucher-Scholz G, et al. Autophosphorylation of DNA-PKCS regulates its dynamics at DNA double-strand breaks. The Journal of cell biology. 2007;177(2):219–29.

70. DeLong ER, DeLong DM, Clarke-Pearson DL. Comparing the areas under two or more correlated receiver operating characteristic curves: a nonparametric approach. Biometrics. 1988;44(3):837–45.

