## Supplementary Tables and Figures for "BUB1 regulates non-homologous end joining pathway to mediate radioresistance in triple-negative breast cancer"

#### Supplementary Table S1

##### Mutation status of TNBC cell lines

| TNBC Cell Lines | Mutation Status |
| --- | --- |
| **SUM159** | BRCA (Wild-type), HRAS (Mutant), PIK3CA (Mutant), TP53 (Mutant) |
| **MDA-MB-231** | BRAF (Mutant), BRCA (Wild-type), CDKN2A (Mutant), KRAS (Mutant), PIK3CA (Wild-type), TERT (Mutant), TP53 (Mutant) |
| **MDA-MB-468** | BRCA (Wild-type), PTEN (Mutant), RB1 (Mutant), TP53 (Mutant) |
| **BT-549** | BRCA (Wild-type), PTEN (Mutant), RB1 (Mutant), TP53 (Mutant) |
| **T-47D** | BRCA (Wild-type), PIK3CA (Mutant), TP53 (Mutant) |

### Supplementary Table S2

#### gRNA sequences for BUB1 knockout

| Guide | Sequence | Exon |
|---|---|---|
| 1 | AGCCCACATGCAGAGCTACA | 2 |
| 2 | TTTACTAGAACATTTAATGA | 3 |

#### Primers for PCR amplification of BUB1-edited section

| Primer | Sequence |
|---|---|
| Forward | TGACATTGGGGTTCCGTGAG |
| Reverse | TCACTAGTGGCACAGAAGTCT |

#### Primer for Sanger sequencing

| Primer | Sequence |
|---|---|
| Forward | TGACATTGGGGTTCCGTGAG |

#### Supplementary Table S3

**List of antibodies used for Western Blotting/Immunohistochemical /Immunofluorescence studies**

| Antibody | Company | Catalog Number | Dilutions Used |
|---|---|---|---|
| Anti-rabbit IgG, HRP-linked Secondary | Cell Signaling | 7074s | 1:10,000 |
| Anti-mouse IgG, HRP-linked Secondary | Cell Signaling | 7076s | 1:10,000 |
| Anti-Ku70 Rabbit mAb | Cell Signaling | 4104S | 1:1000 |
| Anti-Ku80 Rabbit mAb | Cell Signaling | 2180s | 1:1000 |
| DNA-PKcs Phospho antibody (S2056) (E9J4G) Rabbit mAb | Cell Signaling | 68716 | 1:1000 |
| DNA-PKcs Antibody Rabbit mAb | Cell Signaling | 4602s | 1:1000 |
| Phospho-ATM (Ser1981) Rabbit mAb | Cell Signaling | 13050S | 1:1000 |
| ATM (D2E2) Rabbit mAb | Cell Signaling | 2873S | 1:1000 |
| Phospho-KAP-1 (Ser824) Rabbit mAb | Cell Signaling | 4127S | 1:1000 |
| KAP-1 Antibody Rabbit mAb | Cell Signaling | 4123S | 1:1000 |
| Lamin B1 (D4Q4Z) Rabbit mAb | Cell Signaling | 12586S | 1:1000 |
| Recombinant anti-BUB1 Rabbit mAb | Abcam | ab195268 | 1:1000 |
| B-Actin Mouse mAb (HRP Conjugate) | Cell Signaling | 12262 | 1:30000 |
| Ki67 antigen (Dako Omnis) Clone MIB-1 Mouse monoclonal anti-human antibody | Agilent | GA626 | Ready-to-use |
| Anti-phospho-Histone H2A.X (Ser139) | Millipore | 05-636-I | 1:2000 |
| Goat anti-Mouse, Alexa Fluor 488 | Thermo Fisher Scientific | A-11001 | 1:2000 |

#### Supplementary Table S4

##### Primer sequences used in quantitative PCR (qPCR) analysis

| Target Gene | Primer | Sequence (5'-3') | Tm⁰ |
|---|---|---|---|
| GAPDH | Forward | TCGGAGTCAACGGATTTG | 62.4 |
| | Reverse | CAACAATATCCACTTTACCAGAG | 59.1 |
| BUB1 | Forward | GATCGATTACTTTGGGGTTG | 60.8 |
| | Reverse | AAAAAGACCTTCAGGCTTAC | 56.6 |
| BAX | Forward | TCTGAGCAGATCATGAAGAC | 58 |
| | Reverse | TCCATGTTACTGTCCAGTTC | 57.5 |
| BCL2 | Forward | GATTGTGGCCTTCTTTGAG | 59.8 |
| | Reverse | GTTCCACAAAGGCATCC | 59 |
| PCNA | Forward | CTGTGTAGTAAAGATGCCTTC | 55.6 |
| | Reverse | TCTCTATGGTAACAGCTTCC | 56 |
| PRKDC | Forward | GATCTGAAGAGATATGCTGTG | 56.4 |
| | Reverse | GTTTCAGAAAGGATTCCAGG | 60.1 |
| H2AFX | Forward | AATCCAAGCACCTAGATACC | 57.2 |
| | Reverse | CAGAATTCCAGTTCAGAAGC | 59 |
| XRCC5 | Forward | CAGTGAGAGTCTGAGAAAAC | 53.8 |
| | Reverse | TAGGCTGCAATCCTTATAGAC | 57.6 |
| XRCC6 | Forward | AAGAAGAGTTGGATGACCAG | 58.3 |
| | Reverse | GTCACTTCTGTATGTGAAGC | 54.6 |
| CASP3 | Forward | AAAGCACTGGAATGACATC | 57.6 |
| | Reverse | CGCATCAATTCCACAATTTC | 62.8 |
| CASP9 | Forward | CTCTACTTTCCCAGGTTTTG | 57.9 |
| | Reverse | TTTCACCGAAACAGCATTAG | 60.3 |

### Supplementary Fig. S5

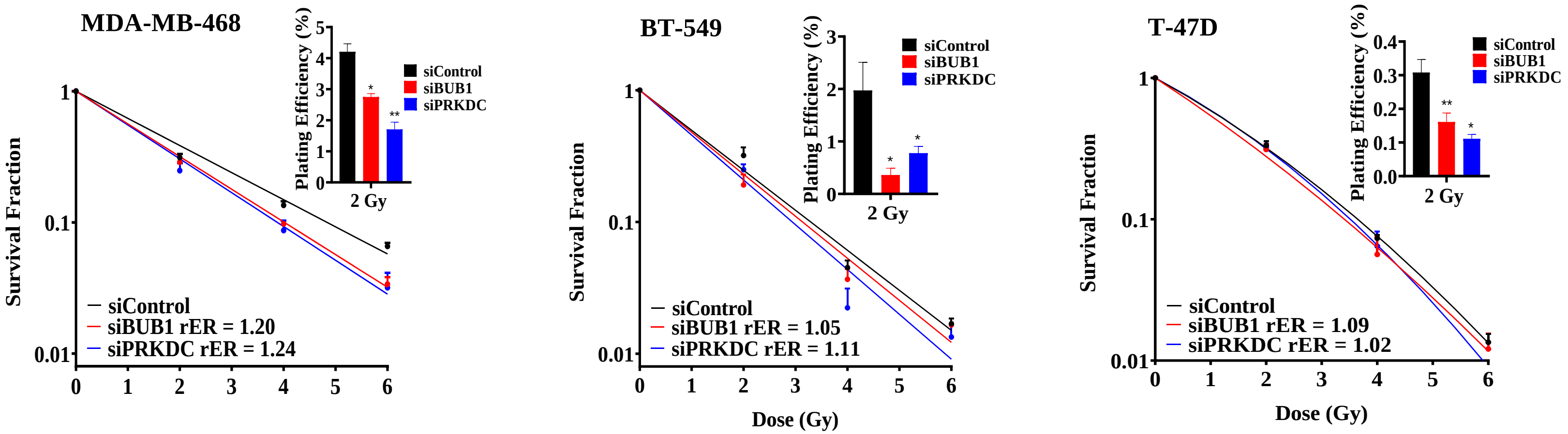

#### Supplementary Fig. S6

**(a)**

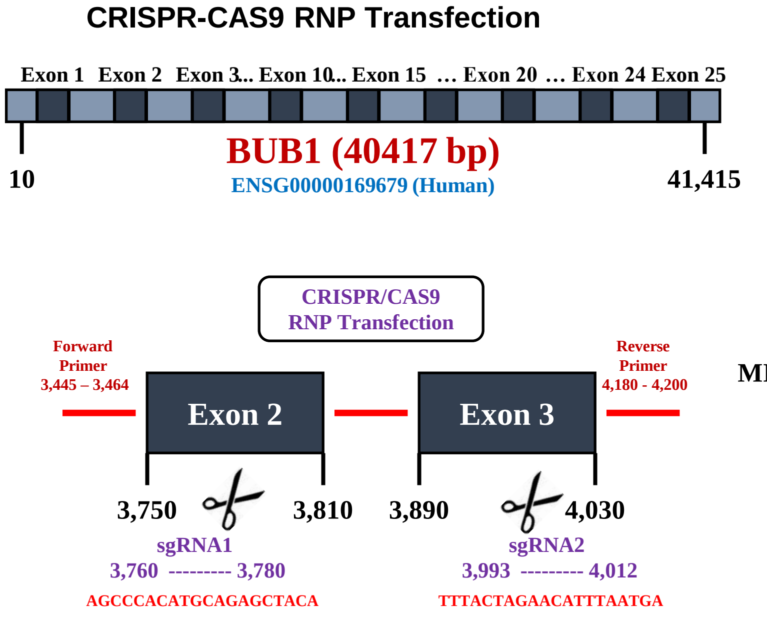


**(b)**

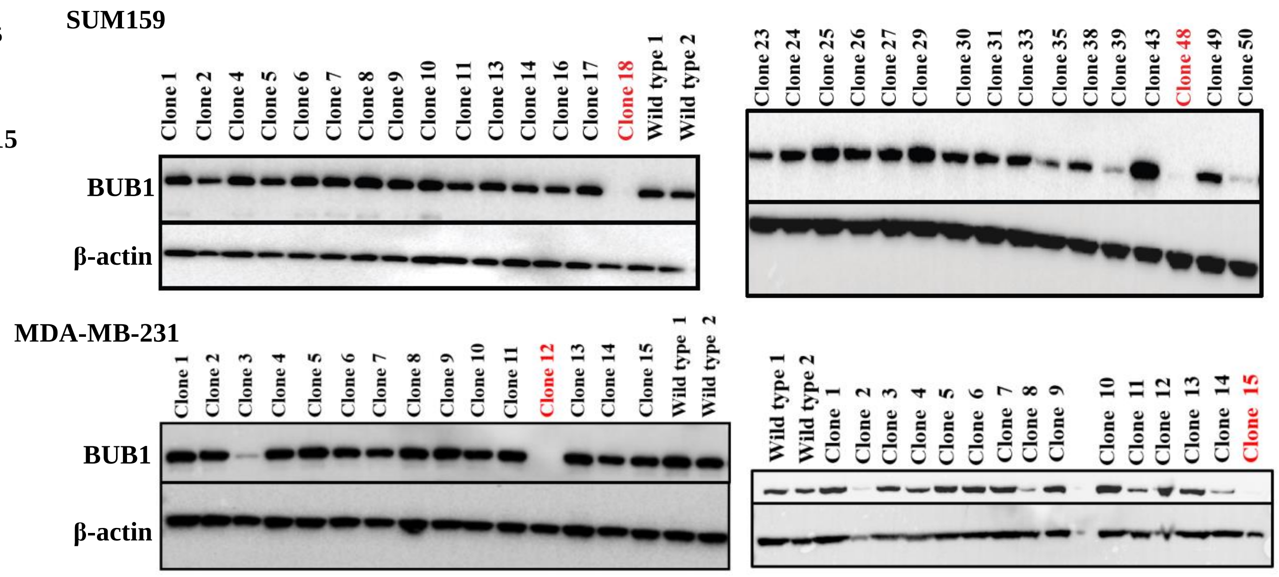


**(c)**

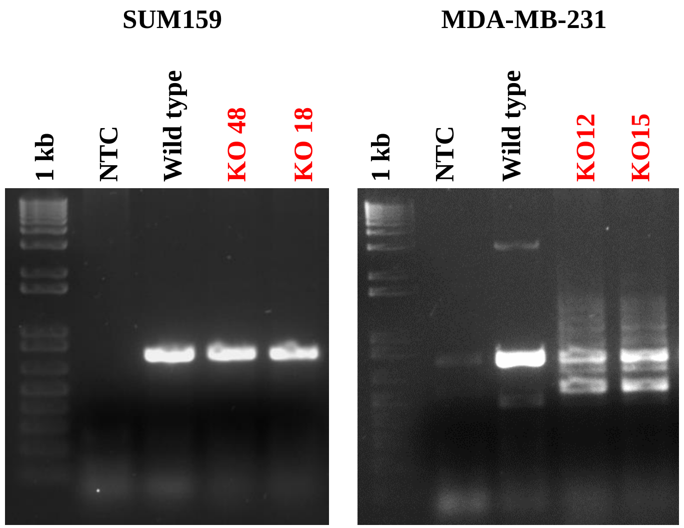


**(d)**

**SUM159 KO48**

**Exon 2**

(WT) AGCCCACATGCAGAGCTACAAGGG
(KO) AGCCCACATGCAGAGCTAT___GGG

**Exon 3**

(WT) TACTAGAACATTTAATGAAGGAATTTTT
(KO) TTTACTAGAACATTTAATGAAGGAATTT

**SUM159 KO18**

**Exon 2**

(WT) AGCCCACATGCAGAGCTACAAGGG
(KO) AGCCCACATGCAGAGCTACGCCAGG

**Exon 3**

(WT) TTTACTAGAACATTTAATGAAGGAATTT
(KO) CTACGACATTNCTTTAAGANAACATTAA

**MDA-MB-231 KO12**

**Exon 2**

(WT) AGCCCACATGCAGAGCTACAAGGG
(KO) AGCCCACATGCAGAGCTATGACCC

**Exon 3**

(WT) TTTACTAGAACATTTAATGAAGG
(KO) CATTCTCTGATTCAACAGACTTC

**MDA-MB-231 KO15**

**Exon 2**

(WT) AGCCCACATGCAGAGCTACAAGGGCAATGA
(KO) AGCCCACATGCAGAGCTATGACCCAAGATT

**Exon 3**

(WT) TTTACTAGAACATTTAATGAAGGA
(KO) GGAAAATCTGTTCATTCTCTGATT

(e)

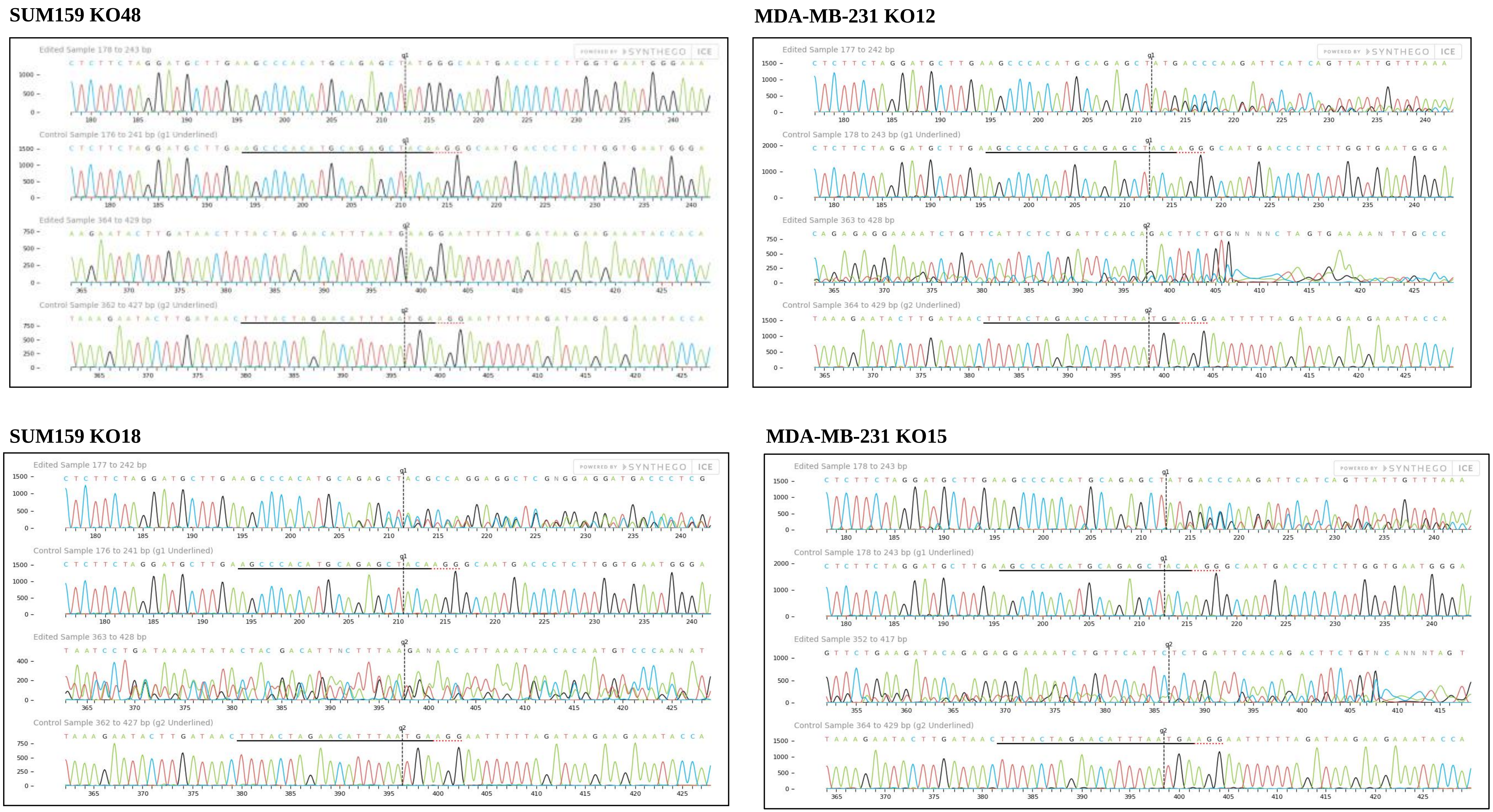
SUM159 KO48
Edited Sample 178 to 243 bp
Control Sample 176 to 241 bp (g1 Underlined)
Edited Sample 364 to 429 bp
Control Sample 362 to 427 bp (g2 Underlined)
MDA-MB-231 KO12
Edited Sample 177 to 242 bp
Control Sample 178 to 243 bp (g1 Underlined)
Edited Sample 363 to 428 bp
Control Sample 364 to 429 bp (g2 Underlined)
SUM159 KO18
Edited Sample 177 to 242 bp
Control Sample 176 to 241 bp (g1 Underlined)
Edited Sample 363 to 428 bp
Control Sample 362 to 427 bp (g2 Underlined)
MDA-MB-231 KO15
Edited Sample 178 to 243 bp
Control Sample 178 to 243 bp (g1 Underlined)
Edited Sample 352 to 417 bp
Control Sample 364 to 429 bp (g2 Underlined)
POWERED BY SYNTHEGO ICE

Supplementary Fig. S7

(a)

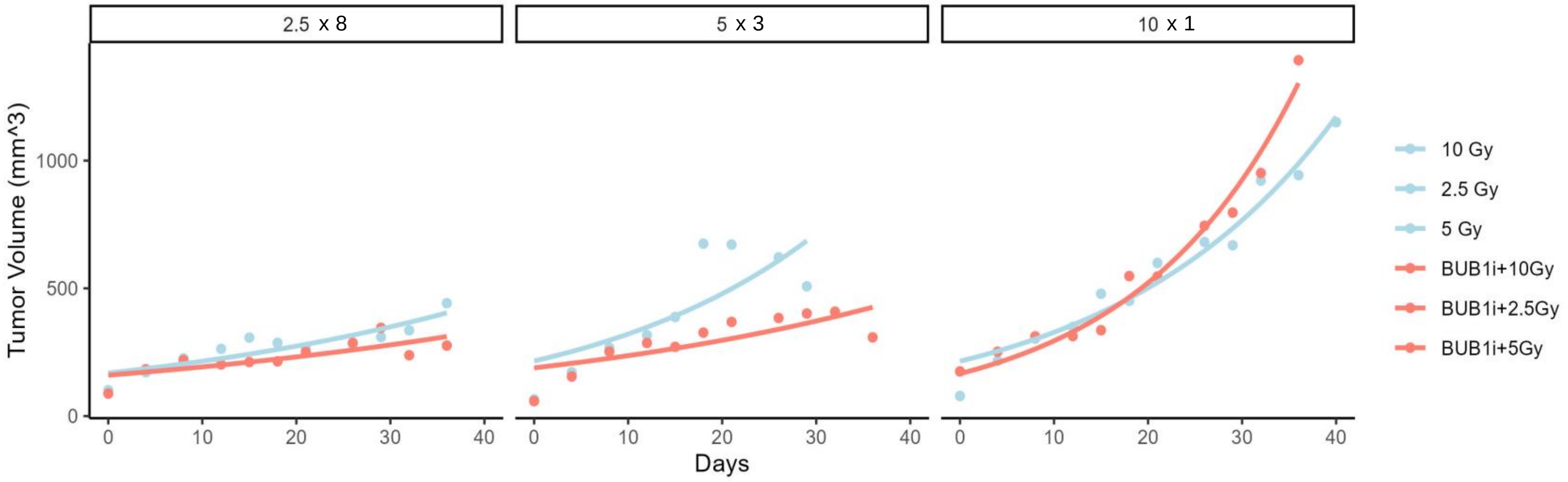


(b)

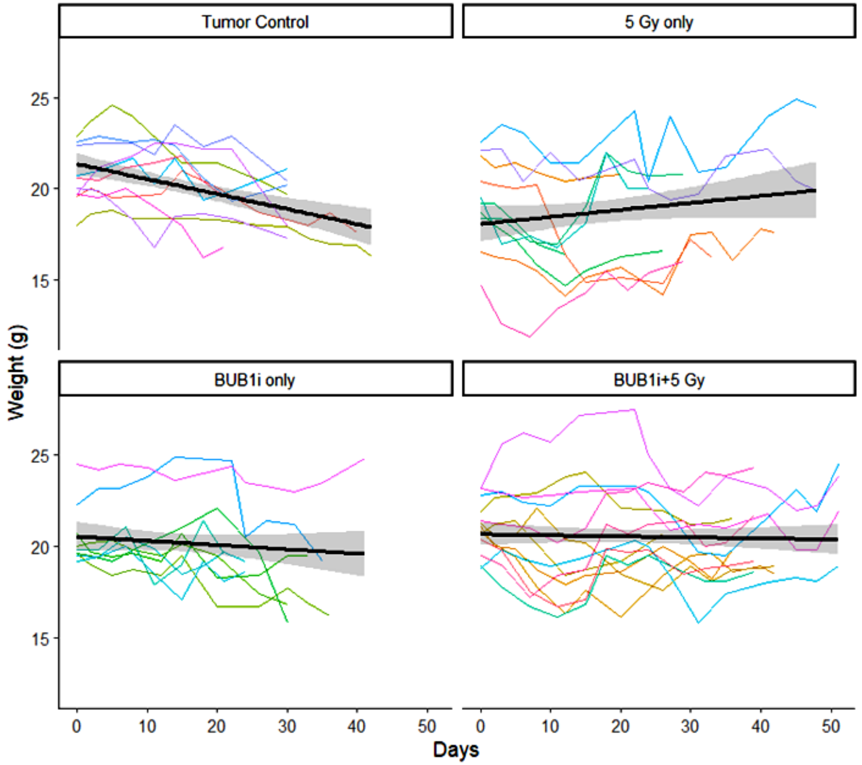

#### Supplementary Fig. S8

(a)

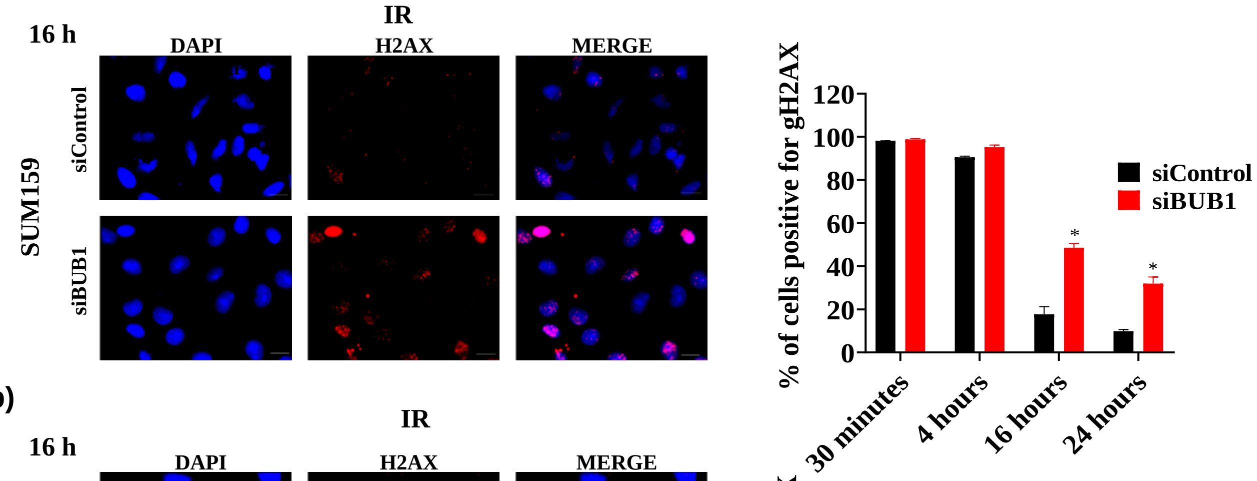


(b)

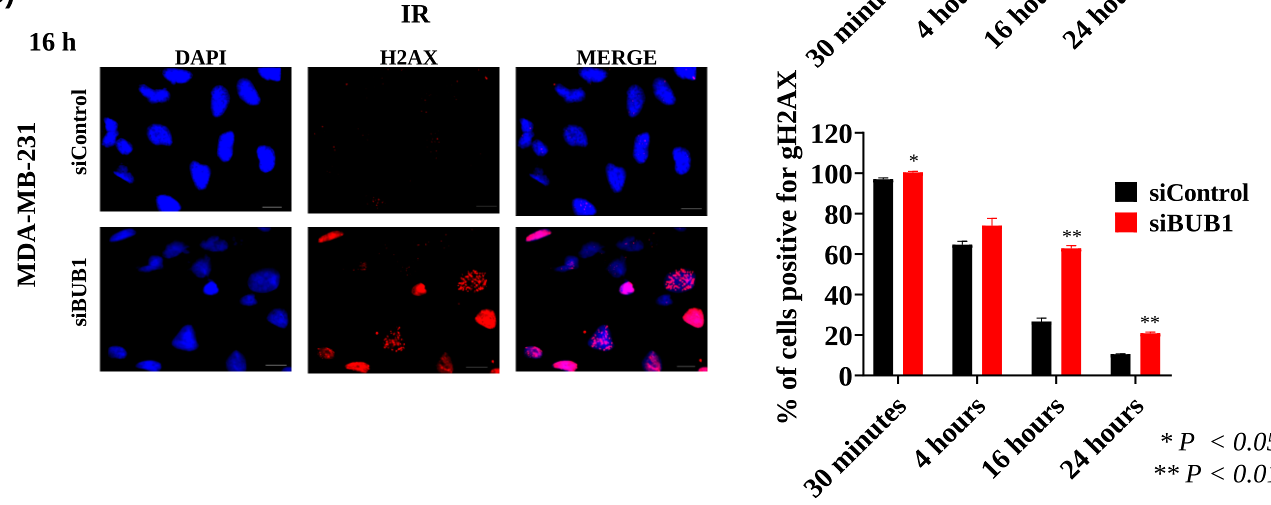


* *P* < 0.05
** *P* < 0.01

#### Supplementary Fig. S9

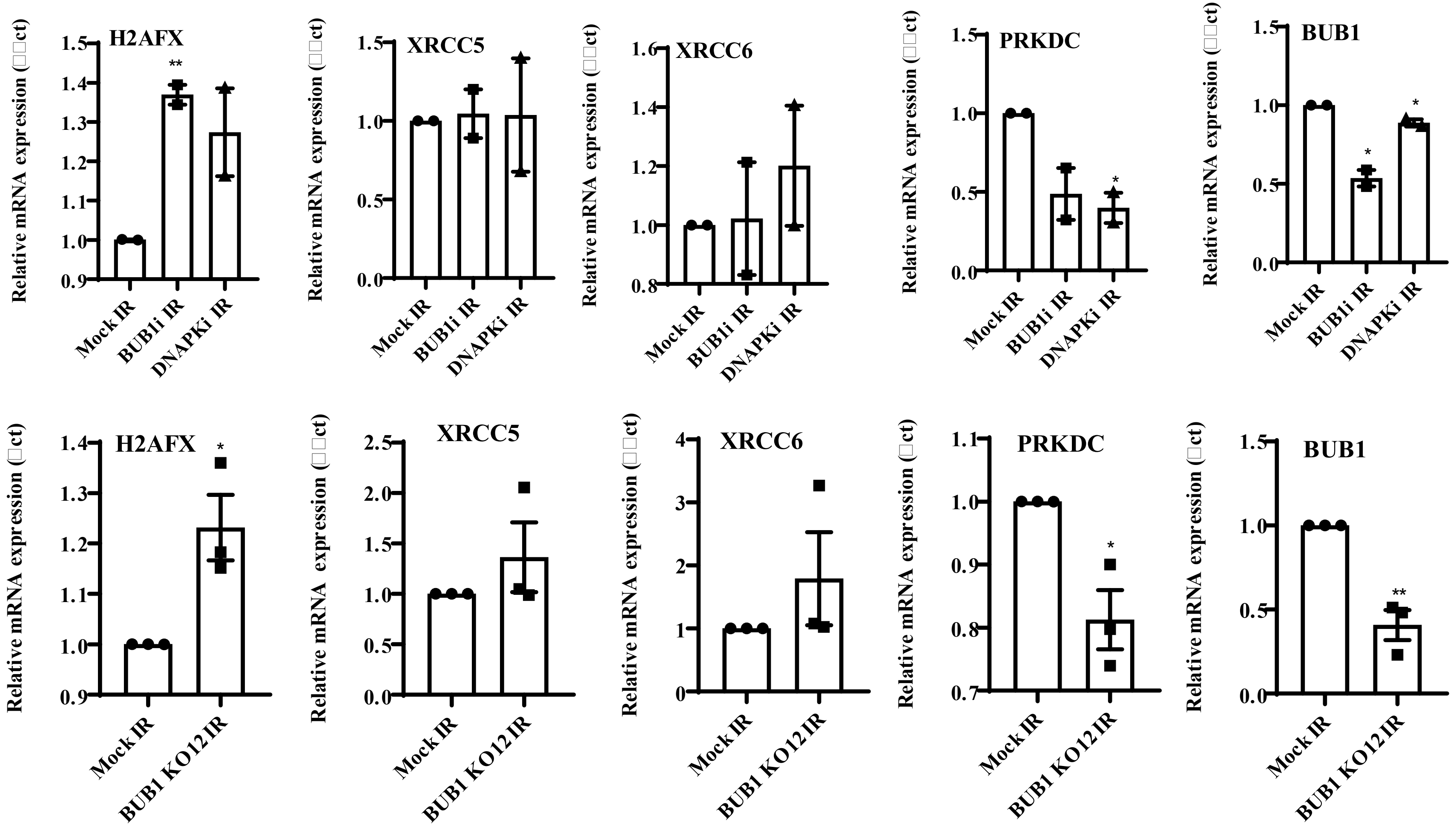

Supplementary Fig. S10

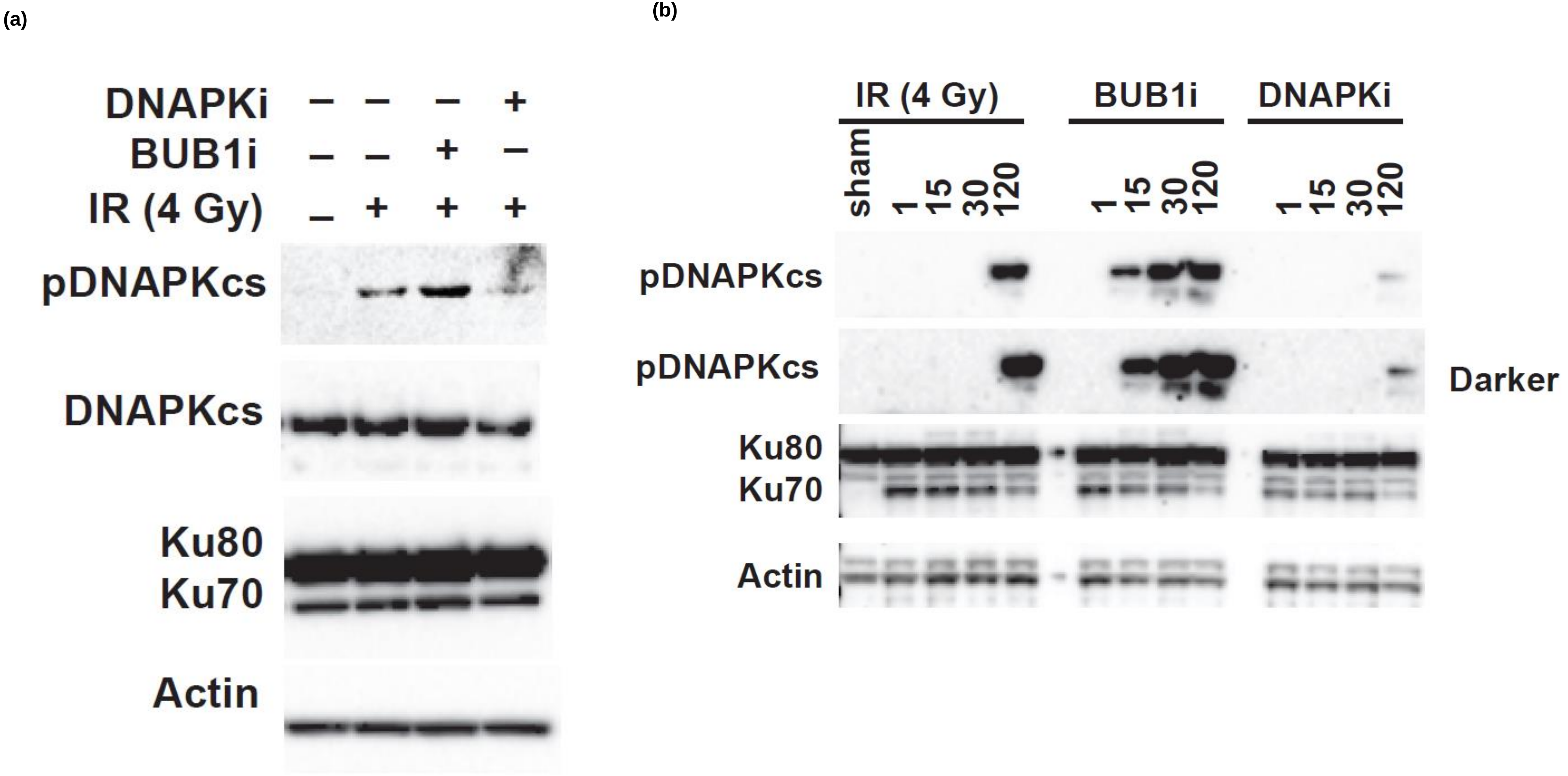

Supplementary Fig. S11

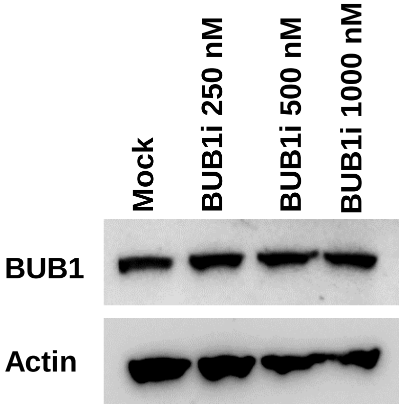

#### Supplementary Fig. S12

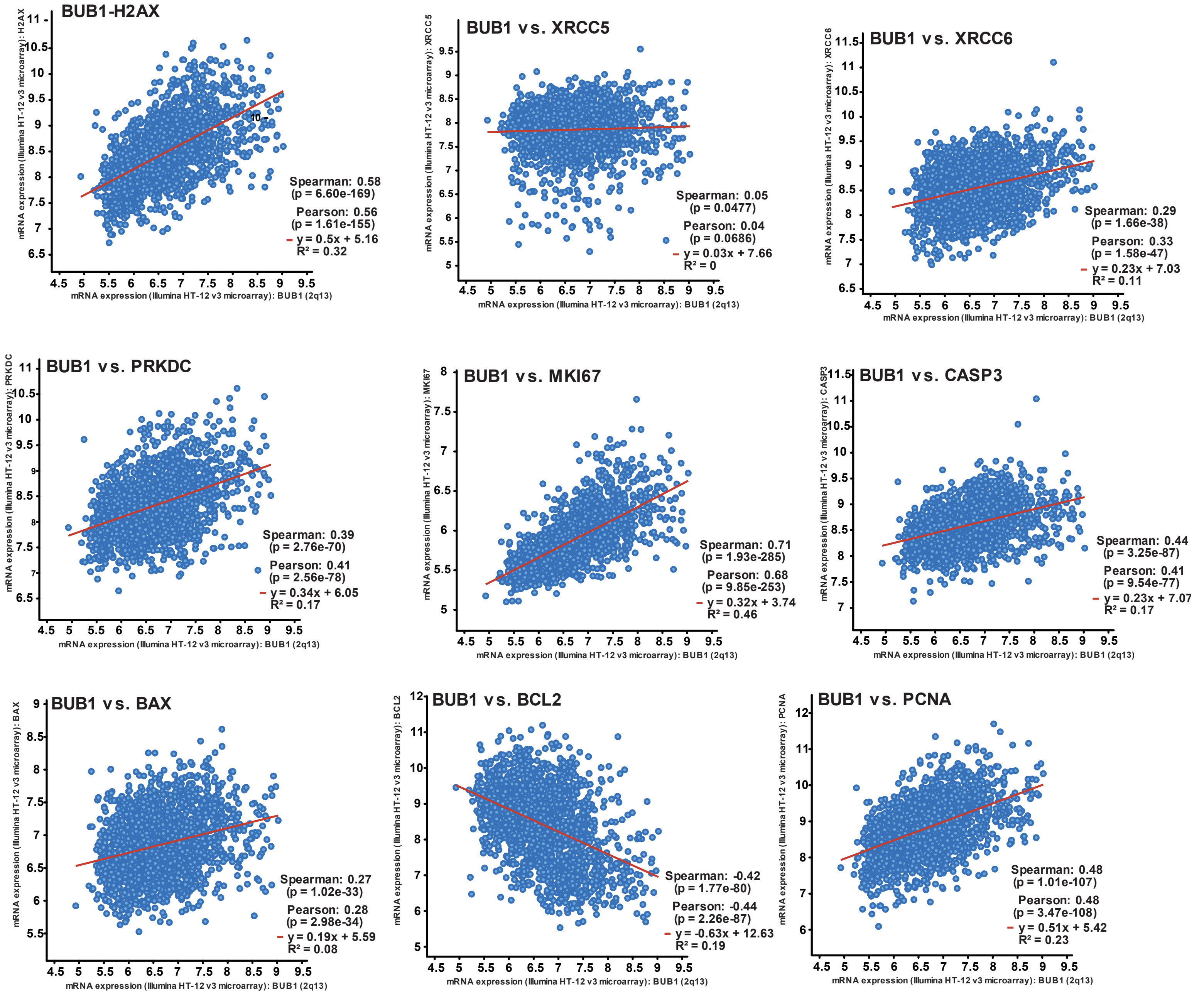

**Supplementary Fig. S13**

**(a)**

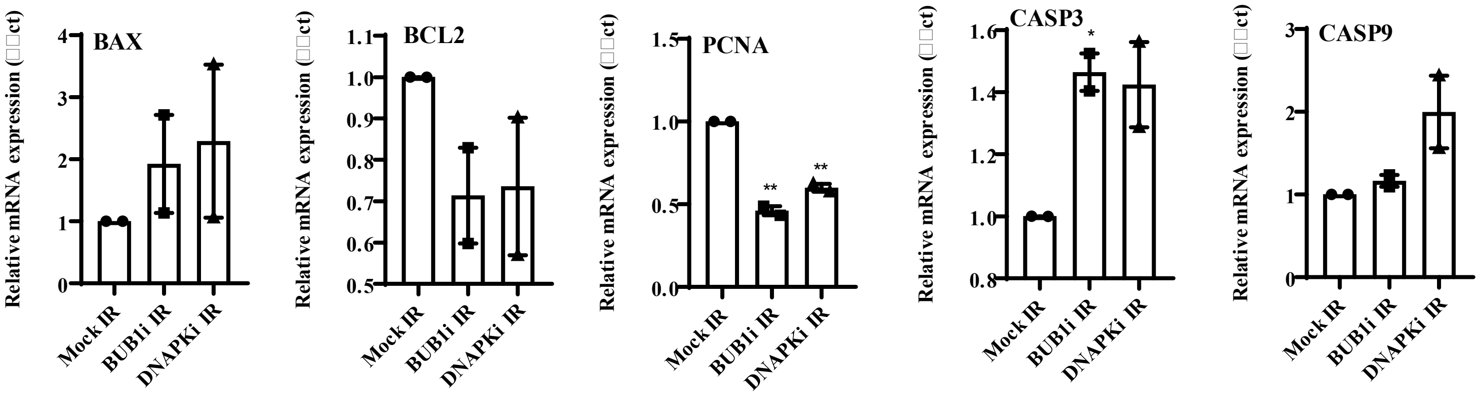


**(b)**

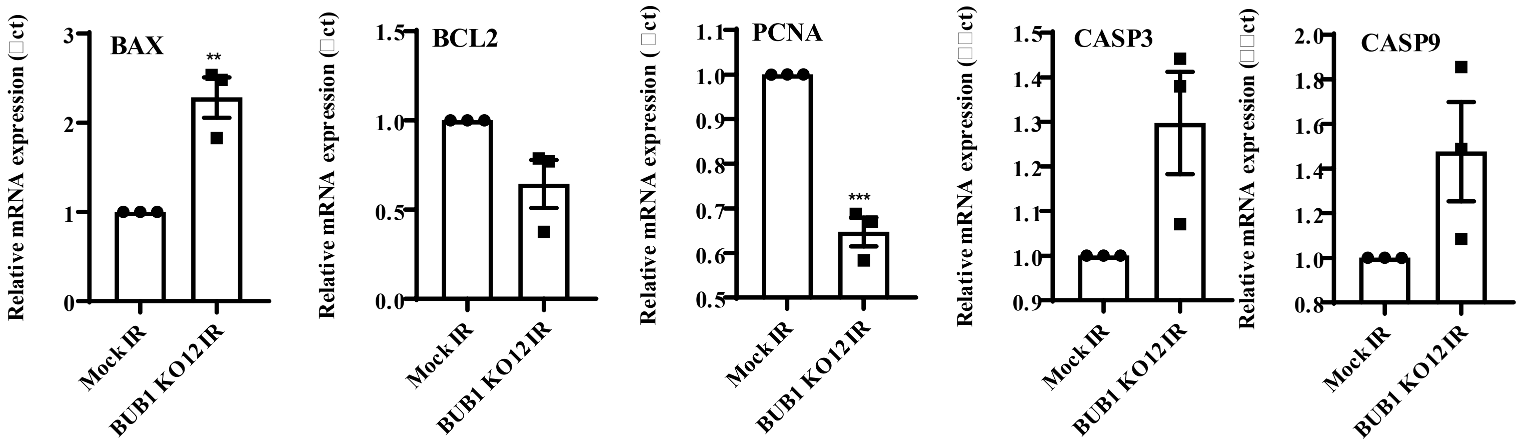
